# Bottom-up and generative computations uniquely explain neural responses across the social brain

**DOI:** 10.64898/2026.02.20.707082

**Authors:** Manasi Malik, Minjae Kim, Tianmin Shu, Shari Liu, Leyla Isik

**Affiliations:** Department of Cognitive Science, Johns Hopkins University, Baltimore, MD 21218, USA; Department of Psychological and Brain Sciences, Johns Hopkins University, Baltimore, MD 21218, USA; Department of Computer Science, Johns Hopkins University, Baltimore, MD 21218, USA

**Keywords:** social perception, theory of mind, generative inverse planning, graph neural networks, fMRI

## Abstract

Making social evaluations from visual input is a core human ability that engages brain regions involved in social perception, including portions of the superior temporal sulcus (STS), as well as higher-level mentalizing regions, such as the temporoparietal junction (TPJ). One common hypothesis proposes that these regions operate hierarchically: social perception regions like posterior STS (pSTS) implement bottom-up computations to generate fast, stimulus-derived representations of social interactions, while mentalizing regions like TPJ perform inverse-planning computations to infer the underlying goals and motivations driving agents’ behavior. However, this computational-neural mapping has never been formally tested, in large part due to the lack of successful computational models of social processing. We developed computational models aligned with these two frameworks: a graph-neural-network (GNN) model that recognizes social interactions by relying on relational visual information, and a generative inverse-planning model that does so by inverting a model of agents’ goals and the physical world. In this preregistered study, we collected fMRI responses while participants watched videos of agentive animated shapes depicting social interactions and compared neural responses to both computational models. Surprisingly, we found that both the GNN and inverse-planning model explained neural responses in pSTS and TPJ, even after controlling for variance explained by the other model. Exploratory analyses, however, revealed a shift from early perceptual processing towards later higher-order reasoning in both regions, suggesting a temporal rather than spatial hierarchy. Overall, this study provides the first evidence that both social perception and mentalizing regions carry out a combination of relational bottom-up and higher-level inferential computations, perhaps on distinct timescales. This work also provides the first comparison of an inverse-planning model to neural activity and demonstrates that theory-driven cognitive models can successfully predict fMRI responses to social scenes.

**Significance Statement:** The ability to recognize social interactions between others is central to humans’ daily lives and engages brain regions supporting social perception and mental state inference. The neural computations underlying this ability, however, are poorly understood. Here we leveraged new models of bottom-up social perception and generative social inference to test the hypothesis that these complementary computations are carried out in separate brain regions. We compared both models to brain responses from subjects viewing procedurally generated videos of social interactions. Surprisingly, we found that both models explained neural activity in both perceptual and mentalizing regions, even when controlling for effects of the other model. These findings challenge the idea of a strict division of labor in the social brain and refine our understanding of the computations supporting human social inference.

## Introduction

The ability to make social inferences from visual input is a core human capacity. Humans can extract rich social meaning from even simple, abstract shapes moving as agents (Heider & Simmel, 1944), including whether their interactions with other agents are cooperative or competitive. This ability emerges early in development, infants demonstrate it within their first year of life (Baillargeon et al., 2016; Geraci et al., 2025; Powell & Spelke, 2018; Spelke, 2022). But what allows the human mind and brain to make such rich inferences from simple visual patterns? While neuroimaging studies have identified a network of brain regions engaged during social inference from visual scenes - including both regions recruited during social perception such as the right posterior superior temporal sulcus (pSTS) and regions recruited during mental state inference such as the right temporoparietal junction (TPJ) - the computational mechanisms enabling these regions to support social inference remain unknown.

Advances in computational modeling now make it possible to test specific hypotheses about these mechanisms. We recently developed two distinct computational models of visual social interaction understanding: a bottom-up model that relationally structures visual information, and a generative inverse-planning model that explicitly simulates agents’ mental states to make social interaction judgments (Malik & Isik, 2023; Netanyahu et al., 2021). Critically, the bottom-up model operates purely on perceptual features without any explicit information about beliefs or goals, while the inverse-planning model explicitly represents and infers latent variables like agents’ goals, beliefs, and relationships by generating rational plans that should match the observed behavior (Baker et al., 2017; Hafri & Firestone, 2021; Papeo, 2020; Ullman et al., 2009; Woo et al., 2022). Our behavioral work showed that both types of computations uniquely explain human social judgments, suggesting that humans employ a combination of both computational strategies (Malik & Isik, 2023). However, the functional organization of these computations in the brain remains unknown.

Previous neuroimaging work has suggested a functional distinction between regions: social perception regions, such as the pSTS, respond selectively to dynamic interactions between agents and contain information about the type of social interaction occurring in a scene but do not represent information about agents’ beliefs (Isik et al., 2017; Walbrin et al., 2018), while mentalizing regions represent more abstract aspects of social cognition, including agents’ goals, beliefs, and intentions (Koster-Hale et al., 2017; Saxe & Kanwisher, 2003; Skerry & Saxe, 2015). One common hypothesis proposes that these regions operate hierarchically, with social perception regions supporting coarse interaction-level representations that are subsequently passed to mentalizing regions for more abstract social reasoning (Ho et al., 2022; McMahon & Isik, 2023). This suggests a specific computational-neural mapping where social perception regions like the pSTS might implement bottom-up relational computations, while mentalizing regions like the TPJ perform inverse-planning computations. For example, observers might first generate a fast, stimulus-derived representation of coarse social interaction information in pSTS (e.g., agents A and B are interacting, or A is chasing B), which could then be refined in TPJ using a slower inverse-planning strategy to infer the underlying goals and motivations driving agents’ behavior (e.g., A intends to tag B in a game versus A wants to harm B)(Gao et al., 2012; Ho et al., 2022). However, this mapping has not been formally tested, largely due to the lack of successful computational models of social interaction recognition, particularly bottom-up models.

In this preregistered study (Malik et al., 2025), we leveraged these two complementary models to test this computational-neural mapping and elucidate the neural computations underlying social scene understanding (Fig. 1). We collected fMRI data from adults while they watched videos of interactions between two animated agents procedurally generated to depict real-life social interactions of cooperation and competition (Netanyahu et al., 2021). Using representational similarity analysis (RSA) (Kriegeskorte et al., 2008), we examined the relationship between neural representations, human behavioral judgments, and two computational models of social interaction recognition: SocialGNN, a bottom-up graph neural network based solely on visual information (Malik & Isik, 2023), and SIMPLE, a generative inverse-planning model based on mental state inference (Netanyahu et al., 2021). We also compared these models to two baseline models: a motion energy model (Adelson & Bergen, 1985) and ControlRNN, a bottom-up recurrent neural network that is matched to SocialGNN but lacks relational inductive biases (Malik & Isik, 2023). We hypothesized that the relational bottom-up model, SocialGNN, would show greater representational similarity than all other models to social perceptual areas such as the pSTS, whereas the inverse-planning model, SIMPLE, would align more closely than other models with areas associated with mental state inference and theory-of-mind, such as the TPJ.

**Fig. 1:**
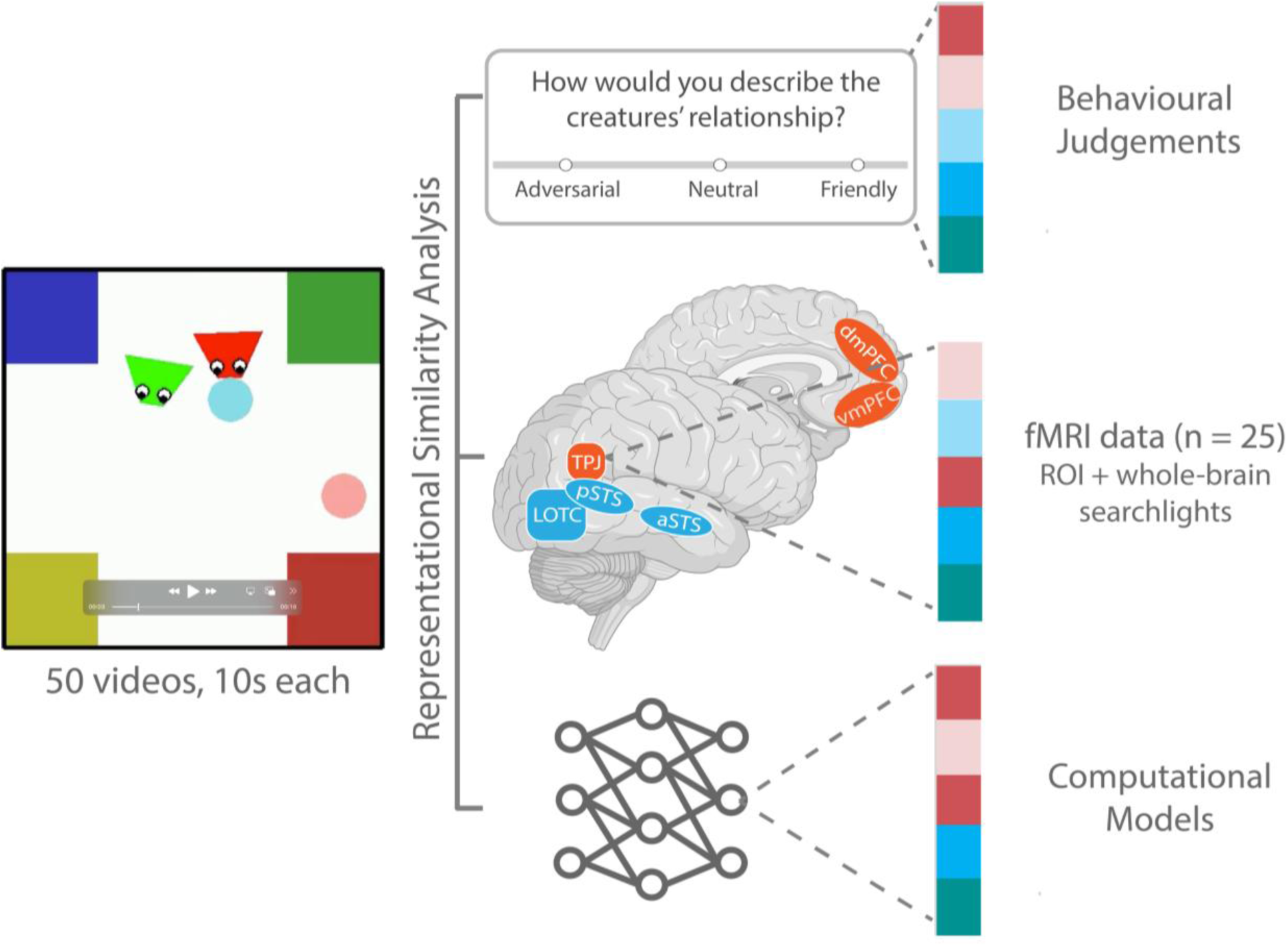
Experimental Overview. Participants (n=25) watched videos of social interactions while their whole-brain fMRI activity was recorded. Neural representations of these videos were then compared to representations from behavioral judgements (e.g., whether the two agents are friendly, neutral, or adversarial) and different computational models (see Fig. 2) in both functionally localized ROIs and whole-brain searchlight analysis.

In both whole-brain and functional region of interest (fROI) analyses, we found that human social judgments significantly explained neural responses across social brain regions, even after controlling for low-level visual information. Surprisingly however, both SocialGNN and SIMPLE significantly explained neural responses in social perception regions (pSTS) *<u>and</u>*mentalizing regions (TPJ), with each model explaining unique variance in both regions. These findings challenge the hypothesis of strict computational segregation across regions and suggests that both regions support a combination of bottom-up relational processing and higher-order inferential reasoning. Exploratory time-resolved analyses, however, revealed that these computations may unfold on different timescales, with an early-to-late shift from bottom-up visual processing toward higher-order inferential reasoning, suggesting a temporal rather than spatial hierarchy. These findings refine our understanding of how the social brain transforms visual input into rich social inferences.

## Results

### 1. Human social ratings, but not motion energy, correlate with social brain regions

Participants (n=25) viewed 10-second videos of animated shapes depicting different social scenes while their neural activity was recorded with whole-brain fMRI (Fig. 1). Videos were selected from the PHASE dataset (Fig. 2a) (Netanyahu et al., 2021) to include the three relationship categories in the dataset, friendly, neutral, and adversarial, and to minimize correlation between our candidate models (see Methods). Participants also completed a series of functional localizers to identify social interaction perception and theory-of-mind regions of interest (ROIs).

**Fig. 2:**
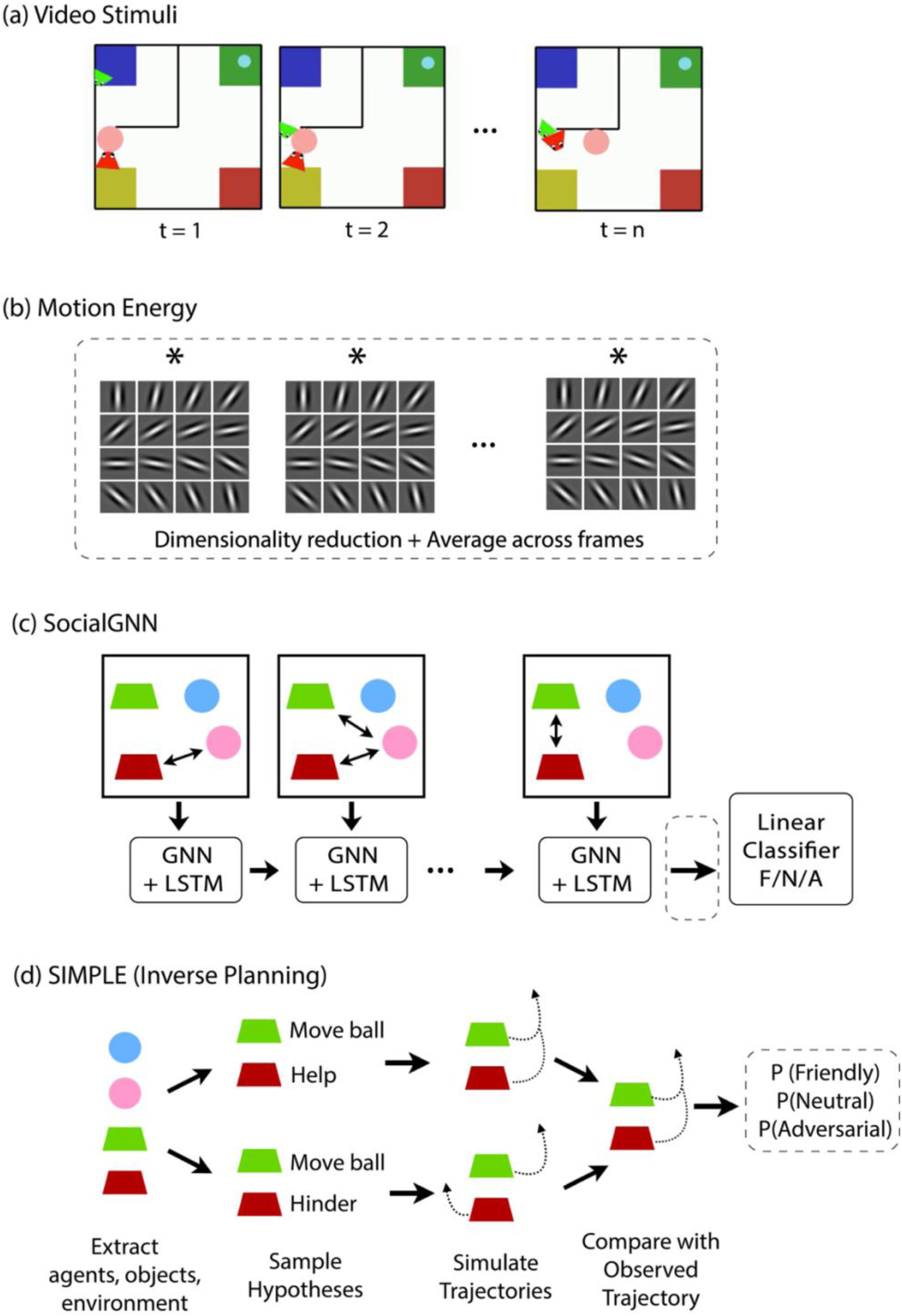
Model Architecture and Representation Diagrams. (a) Example frames from the PHASE dataset videos, depicting two agents moving around in a simple 2D environment, depicting real-life social interactions (b) Motion energy features were extracted from each video by convolving individual frames with a bank of spatiotemporal Gabor filters. These features were dimensionality reduced using principal component analysis (PCA) and averaged across frames to yield a single motion energy representation per video (circled with dotted lines). (c) SocialGNN processes each video frame by constructing a visual graph in which nodes represent agents and objects, with node features including position, velocity, orientation, and agent/object identity, and edges indicating physical contact. The graph is processed by a GNN at each timestep, and temporal information is integrated across frames using a long short-term memory (LSTM) network. The final LSTM hidden state is passed to a linear classifier to predict the relationship category (“friendly”, “neutral”, or “adversarial”) and is used as the SocialGNN representation for each video (circled with dotted lines). (d) SIMPLE is a generative inverse-planning model that, for each video, samples hypotheses about agents’ physical and social goals, simulates corresponding trajectories using a hierarchical planner and a physics engine, and compares these simulated trajectories to the observed trajectories. Based on this comparison, the model assigns probabilities to each relationship hypothesis. The resulting relationship probability vector for each video is used as the SIMPLE representation (circled with dotted lines).

We first asked whether the representational structure of human social judgments matched the representational structure in social brain regions while humans watched these videos. We used human social judgments from our prior experiment (with at least 10 raters per video) (Malik & Isik, 2023) and created a representational dissimilarity matrix (RDM) based on the normalized counts for each rating category ("friendly," "neutral," and "adversarial"). Parallelly, we created RDMs for neural responses in each fROI and for whole-brain searchlights centered on each voxel within a reliability mask. We found that human ratings of social interactions significantly correlated with neural representations in both social perception regions (particularly right pSTS) and mentalizing regions (particularly right TPJ) (Figs. 3a, 4), suggesting that these regions are sensitive to the friendly-neutral-adversarial categorization that people perceive in the videos. Similar results were seen when we used each individual’s post-scan behavioural ratings (Fig. S1a). Specifically, human ratings significantly correlated with bilateral pSTS, TPJ, middle temporal region (MT), and early visual cortex (EVC) (Fig. S1a). Split-half RSA-based reliability of the neural patterns in anterior STS (aSTS) and medial prefrontal cortex (mPFC) regions were not significantly above chance so RSA results for these regions are not reported here and they were excluded from subsequent analyses (Fig. S1b).

**Fig. 3:**
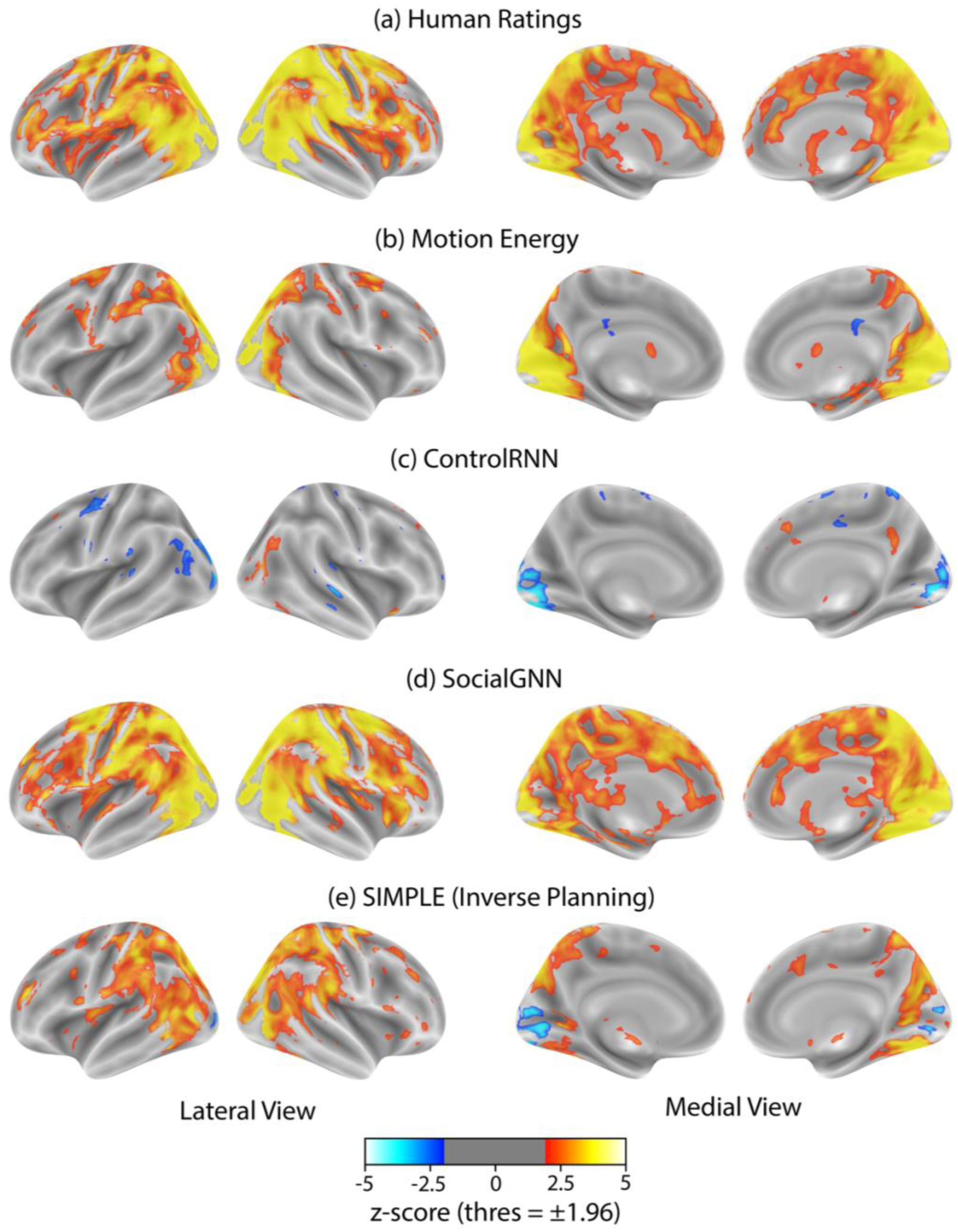
Representational Similarity Analysis: Whole-brain searchlights. Group-level z-scored correlation maps between neural RDMs from whole-brain searchlights and RDMs from: (a) human ratings of social interactions, (b) motion energy, (c) ControlRNN (non-relational bottom-up model), (d) SocialGNN (relational bottom-up model), and (e) SIMPLE (generative inverse-planning model). Z-scores were calculated using two-tailed signed permutation testing with FDR correction across voxels (α = 0.05). Maps were thresholded at ±1.96 to show only significant voxels.

**Fig. 4:**
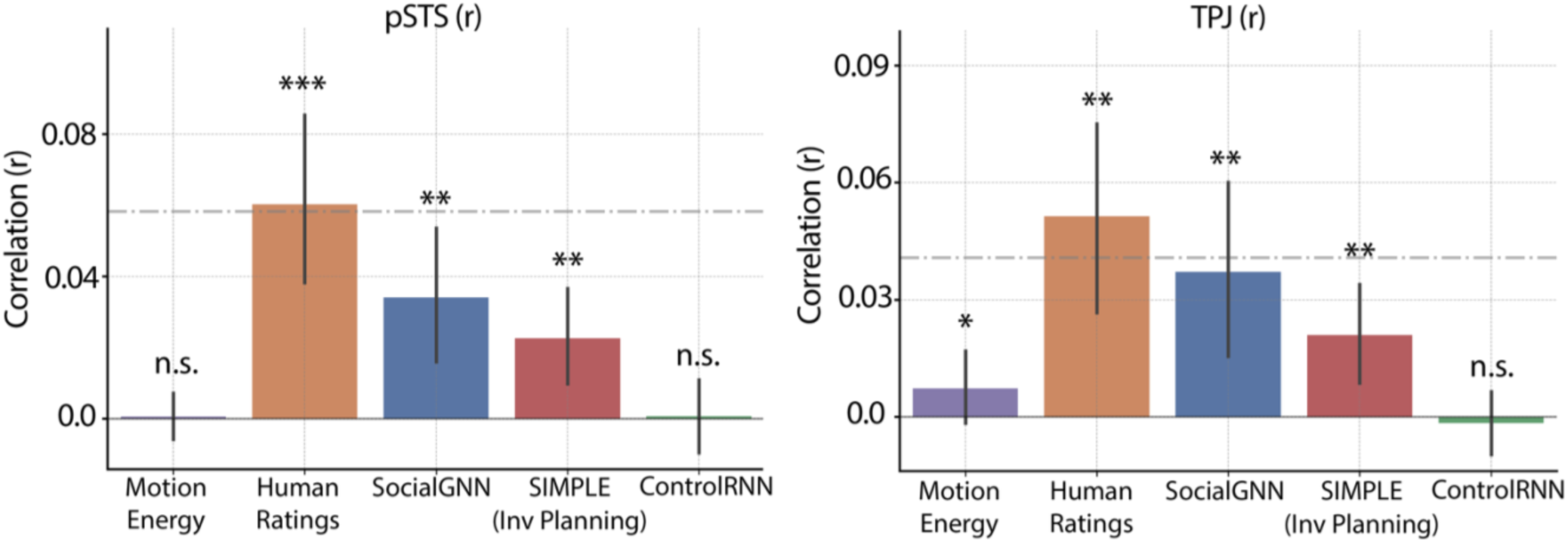
Representational Similarity Analysis: ROI-based. Bar height indicates the mean correlation (across participants), with error bars representing 95 % confidence intervals. Asterisks denote significance levels (p < 0.001 ***, p < 0.01 **, p < 0.05 *); n.s., not significant, evaluated using one-tailed signed permutation testing with FDR correction (α = 0.05). The gray dashed line shows the mean (across participants) split-half RSA reliability for each ROI, corrected using the Spearman–Brown formula. See Table S1 for significant differences between models.

Notably, we also observed representational similarity between these behavioral ratings and early visual regions. To rule out the possibility that low-level visual information was driving our results in the social regions, we examined correlations between neural responses and a motion energy model (Fig. 2b) and found that the pattern of representational similarity differed across early visual and social brain regions. Right EVC was best explained by motion energy in the videos, whereas social regions - right pSTS and right TPJ - were significantly more correlated with human behavioral ratings than with motion energy (Figs. 3b, 4, S1a, & Table S1). Furthermore, variance partitioning analyses revealed that both right pSTS and right TPJ remained significantly correlated with human ratings even after controlling for variance explained by the motion energy model (Fig. S2) or by the pattern of neural activity in EVC (Fig. S3).

### 2. Both bottom-up and inverse-planning models explain variance in social perception and mentalizing regions not captured by low-level visual models

To test our main hypothesis – that social perception regions like the pSTS implement bottom-up relational computations, while mentalizing regions like the TPJ perform inverse-planning computations - we compared the SocialGNN and SIMPLE models to neural responses. SocialGNN is a graph-neural-network-based model that processes videos as sequences of graphs, where nodes in the graph represent agents with visuospatial features (e.g. position, velocity) and edges capture physical contact (Fig. 2c). Each frame, now represented as a visual graph, is processed by a GNN module, and then passed to an LSTM that integrates information across time, to finally a classifier to predict the social interaction (Malik & Isik, 2023). In contrast, SIMPLE (Fig. 2d) is a generative model that infers social relationships through inverse-planning: the model proposes hypotheses about agents’ physical goals and their relationship, simulates what trajectories would result from those hypotheses, and compares them to the observed trajectories to infer the most likely scenario (Netanyahu et al., 2021). We constructed an RDM for SocialGNN using the output from the final LSTM step as the representation for each video, and an RDM for SIMPLE using the predicted probabilities of each social hypothesis (Fig. 2c, 2d).

Surprisingly, we found that both SocialGNN and SIMPLE showed significant representational similarity to both right pSTS and right TPJ (Figs. 3d-e, 4), with no significant differences between the two models (Table S1). We found largely similar trends in the left hemisphere: both models were also significantly correlated with left pSTS, again with no significant differences between them (Fig. S1a, Table S1). In left TPJ, while there were no significant differences between the models, only SocialGNN showed a significant correlation. Both SocialGNN and SIMPLE were also significantly correlated with the pattern of responses in low-level visual regions, including bilateral MT and right EVC (Fig. S1a). To rule out the possibility that correlations in right pSTS and right TPJ were driven by low-level visual features, we again conducted variance partitioning analyses. Both SocialGNN and SIMPLE remained significantly correlated with right pSTS and right TPJ even after controlling for variance explained by the motion energy model (Fig. S2) or by the pattern of responses in right EVC (Fig. S3).

Next, to test whether the relational structure in SocialGNN was critical for explaining neural responses, we compared it with ControlRNN, a matched visual model without graph structure (Fig. S4) (Malik & Isik, 2023). Prior behavioral work underscored the importance of relationally structuring visual input for matching human judgments of social interactions (Malik & Isik, 2023), raising the question of whether such relational organization is also required to explain neural responses. We found that while ControlRNN significantly explained variance in right MT, it did not explain variance in right pSTS or right TPJ (Figs. 3c, 4) and was significantly less correlated with brain responses than SocialGNN in these ROIs (Table S1), demonstrating that relational processing is necessary for matching social brain responses.

We next asked whether our results were driven by the particular representational formats we chose for each model (the array from the final LSTM step for SocialGNN versus the array of predicted probabilities for each relationship category for SIMPLE), rather than the nature of the computations in these models. To test this, we performed exploratory analyses using alternative representations from both models. First, we repeated our analyses using the classifier outputs for SocialGNN rather than the hidden layer activations, as the classifier outputs may be considered more analogous to the SIMPLE’s output probabilities, and found a similar pattern of results in both right pSTS and right TPJ (Fig. S5). Second, we tested “richer” representations from the SIMPLE model by appending the probabilities over each agent’s physical goals to the social relationship probabilities (SIMPLE computes these jointly to make social relationship judgements). We found no significant differences with the original representation in right pSTS or right TPJ (Fig. S6). We also tested a more “internal” representation from the SIMPLE model – the distances between observed trajectories and the model’s internal simulated trajectories for proposed hypotheses – but these performed significantly worse than the relationship probabilities in both regions (Fig. S6), suggesting that this was likely not a good representation for SIMPLE.

Together, these results suggest that both relational bottom-up and inverse-planning computations are carried out across the social brain, and the results are not driven by low-level visual information or idiosyncrasies of model representation choices.

### 3. Bottom-up and inverse-planning models explain *unique* variance in social perception and mentalizing regions

The previous analyses showed that both SocialGNN and SIMPLE correlated with both right pSTS and right TPJ, but did not reveal whether they captured overlapping or distinct aspects of neural responses. To address this, we computed semi-partial correlations between each model and the neural RDMs while controlling for the other model. We found that SocialGNN and SIMPLE both captured significant unique variance in right pSTS and right TPJ (Fig. 5), indicating that each computational process captures distinct information in neural representations in these regions.

**Fig. 5:**
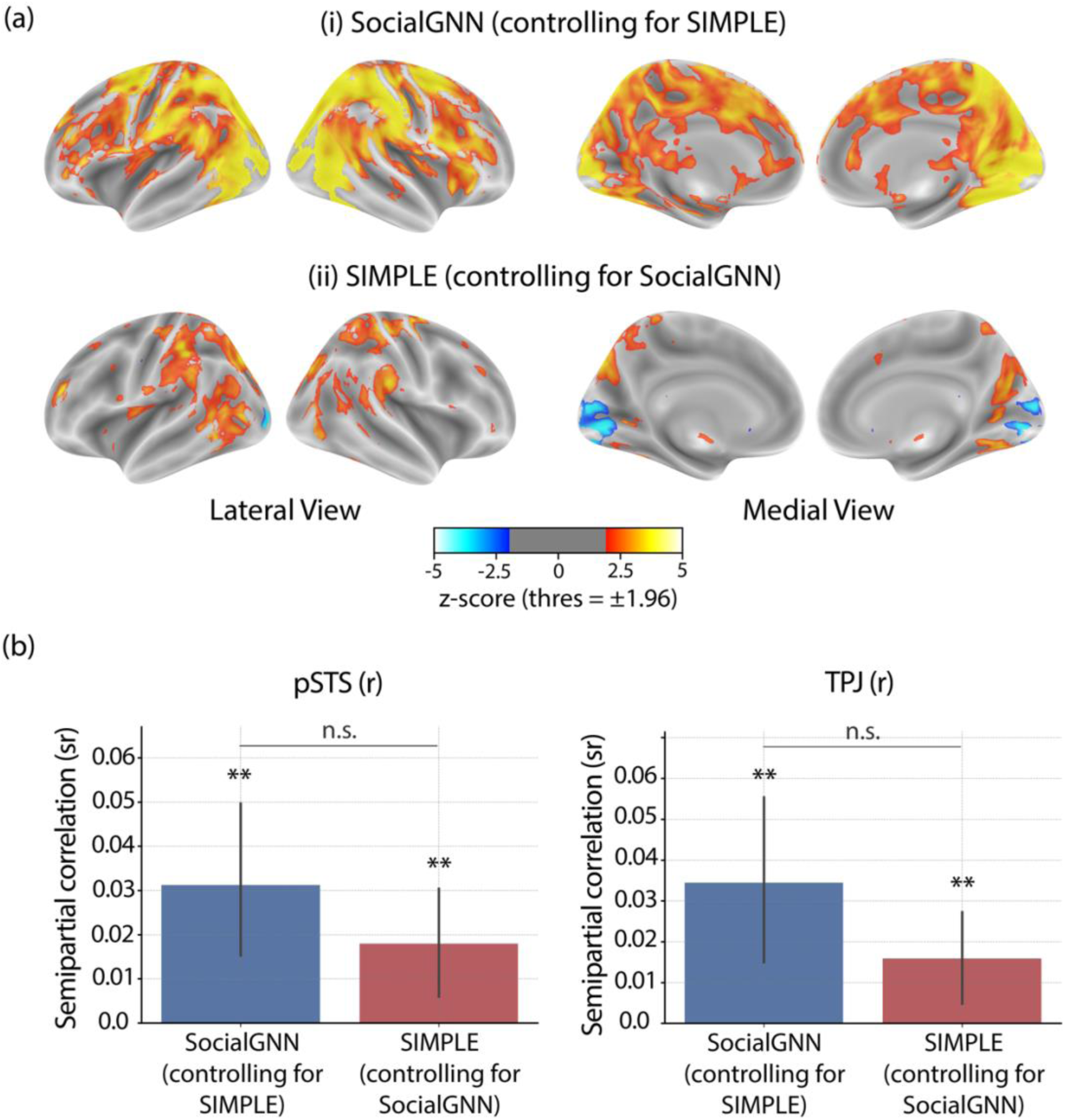
Unique variance explained by SocialGNN and SIMPLE. (a) Whole-brain results (group-level z-scored maps), computed using semi-partial RSA between each model and neural RDMs while controlling for the other model. Statistical significance was determined with two-tailed signed permutation testing and FDR correction across voxels (α = 0.05). (b) ROI-based results. Bar height indicates the mean semi-partial correlation across participants, with error bars representing 95 % confidence intervals. Significance was evaluated using one-tailed signed permutation testing with FDR correction (α = 0.05). Asterisks denote significance levels (p < 0.001 ***, p < 0.01 **, p < 0.05 *); n.s., not significant.

Right pSTS and right TPJ have been shown to be functionally distinct but are anatomically nearby and to some extent overlap in individual participants (Isik et al., 2017; Walbrin et al., 2018). This minimal overlap was also seen in our data (mean DICE coefficient between right pSTS and right TPJ = 0.13, Fig. S8, Tables S2-S3). To see if this anatomical overlap was driving the similar model correlations across regions, we removed overlapping voxels from both ROIs and found that both models continued to explain significant unique variance in both regions (Fig. S7).

### 4. Bottom-up and inverse-planning models show distinct temporal dynamics

We next conducted an exploratory time-resolved analysis to investigate whether the computations in each brain region varied across the course of the 10s videos. Instead of estimating one neural response per voxel for each video, we modeled a separate voxel-wise beta response for each of five consecutive 2s windows. To identify computational differences over time, we computed the semi-partial correlation between SocialGNN or SIMPLE and right pSTS and TPJ responses in each time window, while controlling for the other model.

Interestingly, we found clear differences in the temporal profiles of the two models in both ROIs, with SocialGNN exhibiting a rise-and-fall pattern and SIMPLE showing a more gradual increase over time. Supporting this observation, there was a significant interaction between model and time window in right pSTS (*F(4, 96) = 6.16, p < .001*), indicating that the models’ neural similarity profiles evolved differently over time. This interaction can be seen in the time-resolved profiles with SocialGNN increasing over early windows before declining, and SIMPLE increasing steadily across windows (Fig. 6). In right TPJ, the interaction between model and time window trended toward significance (*F(4, 96) = 2.33, p = .062*), showing a qualitatively similar temporal pattern to that observed in right pSTS. These results suggest that relational bottom-up computations captured by SocialGNN peak during the early-to-middle portions of the video, while higher-level inferential processes captured by SIMPLE build up more gradually over time, in both right pSTS and right TPJ.

**Fig. 6:**
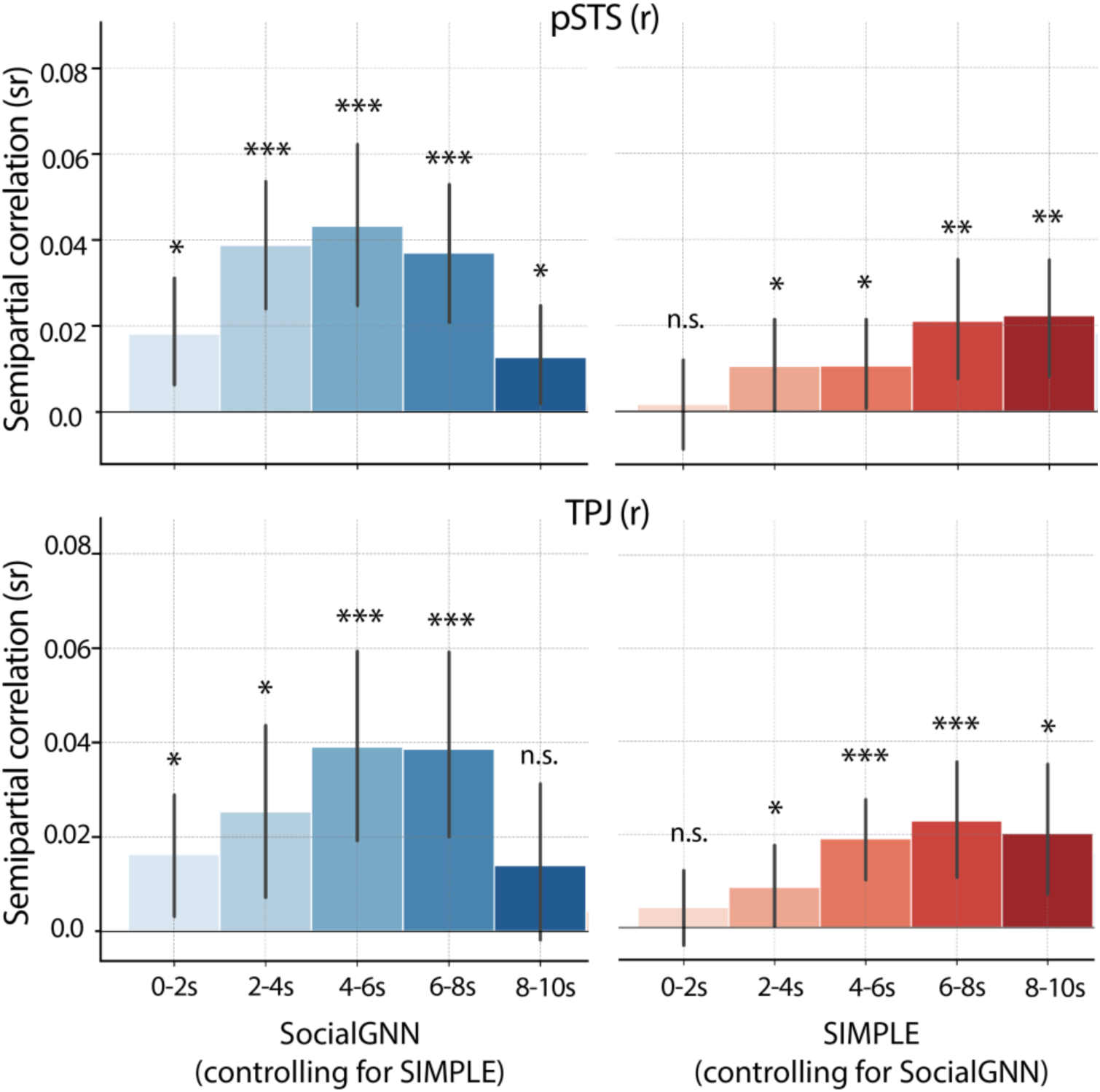
Time-resolved analysis. Semi-partial correlations between each model and neural RDMs in right pSTS and right TPJ, computed using the neural responses in each 2-s window across each 10-s video. Bar heights indicate the mean semi-partial correlation across participants, with error bars representing 95% confidence intervals. Significance was evaluated using two-tailed signed permutation testing with FDR correction (α = 0.05). Asterisks denote significance levels (p < 0.001 ***, p < 0.01 **, p < 0.05 *); n.s., not significant.

## Discussion

In this preregistered study, we used computational models aligned with different theoretical frameworks to investigate the neural computations underlying social scene understanding. We set out to test a previously untested hypothesis: that social perception regions like the pSTS implement bottom-up computations to generate a fast, stimulus-derived representation of social interactions, while mentalizing regions like the TPJ then perform inverse-planning computations to infer the underlying goals and motivations driving agents’ behavior. We leveraged two recently developed computational models of social interaction recognition: a relational bottom-up model, SocialGNN, and a generative inverse-planning model for theory-of-mind reasoning, SIMPLE. Surprisingly, we found that both SocialGNN and SIMPLE showed representational similarity to responses in both social perception regions (particularly right pSTS) and mentalizing regions (particularly right TPJ). These matches in right pSTS and right TPJ were not driven by low-level visual features or by idiosyncrasies in model representations, though relational processing in SocialGNN proved critical for matching neural responses to these social scenes. In follow-up exploratory analyses, we found that both SocialGNN and SIMPLE explained significant *unique* variance in both right pSTS and right TPJ, even after removing overlapping voxels from these ROIs.

Together, our results suggest that there is not a strict feedforward hierarchy in which computations are segregated by region. Instead, both right pSTS and right TPJ perform a combination of relational bottom-up and inverse-planning computations, with both models explaining unique variance in each region. This suggests that SocialGNN and SIMPLE capture complementary rather than redundant computations in both regions.

One important question is whether pSTS and TPJ each independently perform both types of computations, or whether the presence of both computational signatures results from communication between regions that primarily specialize in different computations. Independent processing would suggest greater functional heterogeneity within each region than previously recognized. Alternatively, bidirectional communication - for instance, with relational representations passed from pSTS to TPJ and theory-of-mind inferences passed back from TPJ to pSTS - could appear simultaneous in fMRI despite being sequential, due to fMRI’s poor temporal resolution. Interestingly, our exploratory time-resolved analyses revealed distinct temporal dynamics for the two types of computations. Bottom-up computations showed increasing similarity to neural responses from the beginning to middle of videos before declining, whereas inverse-planning arose later and increased as videos progressed. This temporal pattern suggests a potential shift from early visual processing toward later higher-order reasoning. Since we see similar temporal dynamics in both pSTS and TPJ, these results cannot distinguish whether this temporal progression reflects within-region processing dynamics or rapid communication between regions. Future work using high spatiotemporal neural measurements would be needed to distinguish between these accounts, and would complement other recent research using intracranial human recordings to study social cognition (Jamali et al., 2021).

Beyond the core social brain regions, whole-brain analyses also revealed significant correlation between both computational models and other regions. For instance, both models showed correlations with parietal regions, which could reflect processing of the dynamics and physical interactions in these scenes (Fischer et al., 2016; Pramod et al., 2025). There is growing evidence for interactions between social cognition and physical reasoning systems (Liu et al., 2025). Indeed our stimuli were grounded in physical goals and constraints, and understanding the social interactions likely required appreciating physical aspects of the scene. For instance, determining whether an agent could see a certain object, fit through a given aperture, or had the strength to push/pull objects. In fact, inferring agents’ physical goals and simulating physically plausible trajectories was central to SIMPLE determining their social relationship. While SocialGNN did not make explicit predictions about physical goals, it also incorporated visual features that correlate with the physical world, namely agents’ sizes and wall positions, as graph features. Notably, the distinction between bottom-up and generative simulation we examine here for social cognition has also been asked in the domain of physical reasoning. For example, recent work has compared bottom-up visual prediction models against generative physics simulation approaches (Battaglia et al., 2013; Bear et al., 2020; Bi et al., 2025; Piloto et al., 2022) for physical scene understanding. Testing whether similar computational architectures and neural substrates support both social and physical reasoning could reveal general principles of how the brain transforms visual input into rich conceptual understanding.

It’s important to note that our current analyses test two specific instantiations of bottom-up and generative models, and it is an open question whether our findings would generalize to other models within each framework. Our study already highlights where some simpler bottom-up models (e.g., those without relational processing) fail to match neural responses to social scenes – complementing prior behavioural work (Malik & Isik, 2023). Future work should test whether other graph neural network architectures or alternative models with relational inductive biases would show similar alignment with brain responses. For mentalizing models, while we focused on inverse-planning via SIMPLE, alternative computational approaches to mental state inference exist that do not require physics simulation or full trajectory planning. These include inverse reinforcement learning approaches and causal models that infer mental-states from observed behavior without trajectory simulation (Ho et al., 2022; Jara-Ettinger, 2019), neural models that learn to predict mental states through pattern matching (Rabinowitz et al., 2018), and recent approaches that leverage large-language-models to perform more open-ended inverse planning (Jin et al., 2024; Kim et al., 2025; Shi et al., 2025; Zhang et al., 2026). However, these approaches have primarily been developed in simpler environments (e.g., discrete grid worlds) and/or operate over symbolic state and action representations, rather than continuous, partially observable physical environments like those in our stimuli. Testing whether such approaches adapted to more complex stimuli would show similar alignment with pSTS and TPJ is an important area for future work that would help shed light on mentalizing computations more broadly.

While inverse-planning models have been used to study behavioral judgments in theory-of-mind tasks for decades, to our knowledge, this is the first study to compare the representations from inverse-planning model with neural responses. While we explored different representational choices for SIMPLE, including a combined representation of predicted social relationship and physical goal probabilities (which the model jointly computes to make social relationship judgements), as well as a more "internal" metric based on the model’s internally simulated trajectories for shortlisted hypotheses, more work may be needed to identify the most appropriate “neural” representations to extract from these models. Additionally, while models like SocialGNN can work in image-computable ways (Qin et al., 2025), current inverse planning models are not image-computable since they require an internal world model or simulator and are thus limited to specific types of stimuli. Recent advances in neurosymbolic models that use neural network architecture to do symbolic reasoning seem promising for addressing these limitations (Jha et al., 2024; Yildirim et al., 2020) and may facilitate better model-brain comparisons.

Another promising direction for future work is the development of hybrid models that integrate both bottom-up and top-down processing. Our results suggest that humans employ both types of computations, and an ultimate model of human visual social cognition should combine these mechanisms. A hybrid model could integrate these mechanisms in several ways. For instance, outputs from bottom-up processing could serve as priors for inverse-planning, providing a rapid initial estimate of the interaction type that constrains the hypothesis space for mental state inference. Alternatively, given our current findings of both computational signatures in both regions, a hybrid model might implement bidirectional processing, where bottom-up representations and inverse-planning inferences iteratively refine each other through feedback loops. Such a model might account for the combined variance currently explained separately by SocialGNN and SIMPLE.

Overall, this work represents an important step toward understanding the computational architecture of social scene recognition. Understanding these computations in adult social cognition at the group level, opens the door to future investigations of how these computations might differ across individuals, how they develop, and whether similar or distinct mechanisms are employed in other species (Gandhi et al., 2021; Krupenye & Hare, 2018; Shu et al., 2021; Varrier et al., 2025). The combination of procedurally generated stimuli and cognitive computational models is particularly well-suited for such developmental and comparative investigations, opening new directions in computational social cognition.

## Methods

The experimental design and primary analyses for this study were preregistered prior to data analysis (Malik et al., 2025). The methods described below follow this preregistered analysis plan unless otherwise noted. Any deviations from the preregistered analyses are explicitly identified and justified in the relevant sections.

### 1. fMRI Experiment

All studies detailed here received ethical approval from the Johns Hopkins Homewood Institutional Review Board (IRB) and complied with all relevant ethical regulations. Participants watched visual stimuli and performed tasks inside the fMRI scanner for approximately 2hrs, followed by a post-scan behavioral component.

#### 1.1 Participants

We collected fMRI data from 29 participants (15 Male and 14 Female; mean age = 24.7, range = 18-35yrs; 18 Asian, 3 Hispanic or Latino, 1 Black, 6 White, and 1 Other ethnicities; 27 right-handed and 2 left-handed). All participants had normal or corrected to normal vision, no known neurological diseases or conditions, and complied with MR safety rules. The sample size was chosen based on previous studies where this sample size or smaller has been adequate to reliably detect univariate and multivariate effects in a group analysis (Isik et al., 2017; Zhou et al., 2023). Participants were recruited via flyers in and around the Johns Hopkins Homewood and Medical campuses, and via JHU’s SONA platform where undergraduate students can earn extra credit in select psychology courses by participating in research. All participants provided informed, written consent in compliance with our IRB. Participants were compensated with either $30/hr for 2.5hrs or 3 credits/2.5hrs if they signed up for course credits. 4 participants were excluded from our analyses because they met one or more of our preregistered exclusion criteria: excessive motion during the scans (more than 1 main task run with more than 20% frames with >0.5mm frame displacement); participant not following task instructions during the scan; participant’s post-scan behavioral ratings were 2 standard deviations from the group’s ratings.

#### 1.2 Stimuli

We use the PHASE stimulus set (Netanyahu et al., 2021) which consists of animated shape videos generated via a physical simulator and hierarchical planner, where two agents are moving around in a simple 2D environment, depicting real-life social interactions (Fig. 2a, left). There are also two objects in each video that the agents can push and pull. The environment has four landmarks and some walls, all stationary. The agents and objects (entities) can move over landmarks but cannot pass through the walls. The agents each have a physical goal (e.g. “take pink object to yellow landmark”) or a social goal (e.g. “help the other agent”) that is input to the video generator, and both are given eyes and a triangular body shape to increase the impression of animacy and to show their heading direction. The sizes of the agents and objects, the strength of the agents, and the layout of the walls can all vary across the videos. The dataset includes a 400-video standard dataset and a 100-video generalization set with novel environment layouts and agents with social/physical goals that are unseen in the standard 400-video set. For more details on the dataset see Netanyahu et al., 2021. For this study, we selected 50 videos from the 100-video generalization set – see below for details. We then trimmed these videos to keep only the middle 10s for each in order to have uniform duration videos for the fMRI experiment. Two authors (M.M. and L.I.) independently classified each 10s video as depicting a friendly, adversarial, or neutral interaction, and compared the ratings to the mode behavioral ratings previously collected (Malik & Isik, 2023). For five videos, where author ratings differed from the group mean on full videos, we selected a different 10s segment to better match the group rating. We also used nine videos from the 400-video standard dataset (3 rated as friendly, 3 neutral, and 3 adversarial by the majority of participants) to serve as attention check videos for our experiment. We trimmed these attention-check videos as well to get the middle 10s.

##### Subset selection

We first selected all videos from the PHASE generalization set that were at least 10 seconds long, reducing the set from 100 to 84 videos. To ensure a well-balanced and representative stimulus set, we performed 1,000 iterations of random sampling, selecting 50 videos (without replacement) per iteration from this pool. For each sample, we generated Representational Dissimilarity Matrices (RDMs) for SocialGNN, SIMPLE (Inverse-Planning Model), and Human Ratings and computed their pairwise correlations. Our previous work (Malik & Isik, 2023) found that both SocialGNN and SIMPLE significantly correlated with human judgments on the PHASE dataset but also captured significant unique variance, indicating that they have distinct representations. To reflect this distinction in our fMRI experiment, we selected a final set of 50 videos that met the following criteria: (i) low correlation between SocialGNN and SIMPLE RDMs, (ii) high correlation between SocialGNN and Human Ratings RDMs, (iii) high correlation between SIMPLE and Human Ratings RDMs, and (iv) a balanced distribution of friendly, neutral, and adversarial interactions. The final subset consisted of 16 friendly, 19 adversarial, and 15 neutral videos, with correlation values of r = 0.27 (SocialGNN & SIMPLE), r = 0.50 (SocialGNN & Human Ratings), and r = 0.46 (SIMPLE & Human Ratings), all p < 0.001. This methodology ensures that our stimulus set captures the distinctive computations embodied by SocialGNN and SIMPLE on social scene understanding while maintaining strong alignment with human judgments.

#### 1.3 Experiment Paradigm

The scan protocol for each participant consisted of one high resolution anatomical scan, 10 runs of our main task (‘Main’), 2 runs of a social interaction perception localizer (‘SI-pSTS’), 2 runs of a theory-of-mind localizer (‘ToM’), and 1 run of an intuitive physics localizer (‘Physics’) (Fig. 7). Before going into the scanner, participants were given instructions for each of the four types of tasks they would be doing. Individual tasks are described in detail below. The total scan duration was set to 1hr 33mins. Post the fMRI scan, each participant completed a behavioral task where they watched the same 50 PHASE dataset videos used in the fMRI experiment, outside the scanner. They also completed an Autism Spectrum Quotient (AQ) (Baron-Cohen et al., 2001).

**Fig 7:**
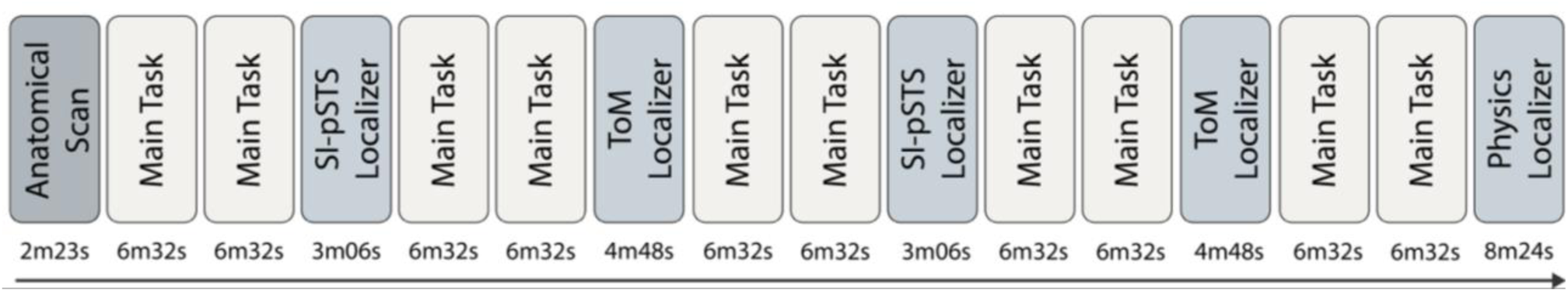
fMRI Experiment Paradigm.

##### Main Task

For our main task, the participants were shown the 50 selected videos from the PHASE dataset. The instructions given to them were: *“You will watch some videos with two creatures moving around. Pay attention to the relationship between the two creatures. After some randomly picked videos, you will be asked to describe their relationship as Friendly, Neutral, or Adversarial. Follow the instructions on the screen to give your response.”* The 50 videos were randomly split into two groups of 25 videos each, and this process was repeated 5 times for each participant, giving us 10 runs. Each video was presented for 10s, followed by a 2s interval where a gray screen was shown. For five randomly selected trials, an additional 2s (1 TR) jitter was added to the interval to improve signal deconvolution in the GLM. In each run, 3 attention check videos were randomly picked from the 9 attention check videos and interleaved with the 25 main videos. After each of these attention-check videos, the participants were a shown a question on the screen “*How would you describe the creatures’ relationship?”* and they had to respond with “friendly”, “neutral”, or “adversarial” using the buttons given to them. The button order varied randomly for each trial, with corresponding labels displayed. Attention check trials were discarded during analyses and there was no question presented after the other videos. Video presentation order was also randomized. The duration for each run of this task was 6mins 32sec.

##### SI-pSTS localizer

In this task, participants passively watched point light figures either interacting (social interactions) or performing independent actions (nonsocial actions) from Isik et al., (2017). The duration of each run in this task was 3mins 6secs.

##### ToM localizer

In this task, participants performed the efficient False Belief Localizer from Dodell-Feder et al., (2011). Participants were asked to read a small paragraph on screen and then answer a True/False question about it. The paragraph-question pair could either be a false-belief condition or a false-photo condition. The duration of each run in this task was 4mins 48s.

##### Physics localizer

In this task, participants completed a task from Fischer et al., (2016) where they watch 10s videos of two dots moving either as physical objects or interacting socially. During a 2s disappearance of one dot, participants are asked to imagine its trajectory and judge whether it reappears where they expected it to. The duration of each run was 8mins 24secs.

##### Post-Scan Behavioral Ratings

After completing the fMRI scan, participants engaged in a post-scan behavioral task that involved watching the same set of 50 videos from the PHASE dataset that were used during the scanning session. This task mirrored the Main Task performed inside the fMRI scanner, except that now the participants were asked to describe the creatures’ relationship after every video. Further, unlike the in-scanner task, this post-scan session did not include attention check videos, and responses were recorded via the arrow keys on a laptop. The behavioral task lasted approximately 11 minutes on average across participants (range ∼ 10–13 minutes).

##### Autism Spectrum Quotient (AQ) Assessment

After the fMRI scan, participants completed the standard Autism Spectrum Quotient (ASQ) questionnaire (Baron-Cohen et al., 2001). It consisted of 50 statements, where participants responded with "Definitely Agree," "Slightly Agree," "Slightly Disagree," or "Definitely Disagree." Responses were scored using standard ASQ scoring methods, with each autistic-trait-consistent response contributing one point, yielding a total score ranging from 0 to 50. Exploratory analyses showed no significant relationship between AQ and the neural measures reported in the main text.

#### 1.4 MRI Data Acquisition Parameters

All fMRI scans were conducted on a 3T Philips Elition RX scanner (with a 32-channel head-coil) at the F.M. Kirby Research Center for Functional Brain Imaging at the Kennedy Krieger Institute. The anatomical scans were performed with a T1-weighted magnetization-prepared rapid-acquisition gradient-echo (MPRAGE) sequence with the following parameters: repetition time (TR) = 7.6 ms, echo time (TE) = 2.4 ms, flip angle = 18°, voxel size = 1 x 1 x 1 mm^3^, field of view = 240 x 240 x 180 mm^3^. The scans were collected with sagittal primary slice direction. The functional data were acquired using a T2∗-weighted gradient-echo echo-planar imaging (EPI) sequence, sensitive to blood oxygen level-dependent (BOLD) contrast, with the following parameters: repetition time (TR) = 2s, echo time (TE) = 30 ms, flip angle = 77°, voxel size = 3 x 3 x 3 mm^3^, field of view = 220 x 220 x 118.5 mm^3^, and 34 axial slices spanning across the entire cortex.

#### 1.5 MRI Data Preprocessing

##### Missing runs

Due to time constraints, the 9th and 10th runs of the Main task (corresponding to the 5th repeat of each video) were not conducted for 3 participants. For these participants, analyses were performed using the first 4 repeats of the Main task, which was sufficient for all reported analyses. In addition, the Physics localizer was not conducted for 1 participant.

##### Converting to BIDS format and defacing the scans

The data was converted to BIDS format, and the anatomical and functional scan files were converted from PAR/REC format to NIFTI format. The anatomical images were defaced using the tool *pydeface* (Gulban et al., 2022).

##### Preprocessing using fMRIprep

Preprocessing was conducted using fMRIPrep version 21.0.2 (Esteban et al., 2018), built on Nipype 1.6.1 (Gorgolewski et al., 2011; K.J. Gorgolewski et al., 2018). Default settings were applied, with output spaces set to T1w (participant native space), MNI152NLin2009cAsym, and fsnative (participant-specific native surface space generated by FreeSurfer).

##### Generalized Linear Modeling of fMRI data

After preprocessing, we took the functional data in MNI152NLin2009cAsym space and smoothened it using a 4mm FWMH kernel implemented in the smooth_img function from nilearn. We then used GLMsingle (Prince et al., 2022) to estimate individual trial responses (betas). We used all three components of GLMsingle: using a library of HRFs to find a potential HRF for each voxel, GLMdenoise (Kay et al., 2013), and fractional ridge regression. We set ‘autoscale’ and ‘percentbold’ as 1. In an exploratory time-resolved analysis, instead of estimating a single response per trial (each trial corresponding to a 10s video), we estimated voxel-wise responses for five consecutive 2s windows spanning each trial, using the same GLMsingle settings described above.

##### Processing estimated betas

Estimated betas were normalized (voxelwise) for each task for a participant and then averaged across repeats per condition (each video stimulus).

### 2. Analyses

To look at the representational similarity between different brain areas and the computational models and behavioral judgements, we performed representational similarity analysis first using whole-brain searchlights and then using participant-specific ROIs.

#### 2.1 Whole-brain Searchlights RSA

For each participant, we defined (overlapping) spherical volumetric searchlights with a radius of 3 voxels across the whole brain. Custom Python code was used to extract neural activity patterns within each searchlight for every condition in the Main task, and representational dissimilarity matrices (RDMs) were computed using 1 - Pearson correlation for each condition pair. These neural RDMs were then compared to computational model RDMs and behavioral data RDMs using Spearman’s rank correlation. To generate group-level maps, we deviated from the preregistered parametric second-level model (Nilearn SecondLevelModel) and instead used a nonparametric two-tailed signed permutation test (5,000 iterations) with FDR correction across voxels (α = 0.05), which provides more reliable inference for correlation-based RSA measures that may violate normality assumptions. The resulting p-values were converted to Z-statistics for visualization. This group-level approach tests for consistent effects across participants with the resulting *z*-scored group maps highlight areas of significant representational similarity. Since all individual analyses were conducted in MNI space, no additional realignment was necessary.

In addition to standard RSA, we also conducted variance-partitioning analyses using semi-partial Spearman correlations to estimate the unique contribution of each computational model or behavioral RDM while controlling for other predictors. Significance testing was performed identically to the procedure described above.

#### 2.2 ROI-based RSA

##### ROI Definitions

For each participant, we defined ROIs using predefined parcels and their activations in the functional localizer tasks (Tables 1, S2; Fig. S8). We generated a contrast map for each localizer task by taking the difference of beta maps for the two conditions: (i) ‘interaction’ - ‘no-interaction’ in the SI-pSTS localizer, (ii) ‘belief’ - ‘photo’ in the ToM localizer, and (iii) ‘physical’ - ‘social’ in the Physics localizer. We then took the top 10% of voxels in the contrast map lying inside the respective parcel, which was converted to binary to obtain the ROI mask. We separated the ROI mask into left (L) and right (R) hemispheres for all ROIs except those in the mPFC. To investigate overlap between nearby ROIs we computed the pairwise DICE coefficient (Table S3). We don’t use the Physics localizer for the analyses presented in this study.

**Table 1:** ROI Definitions are based on the cited papers with the following exceptions. *The STS parcel from Deen et al., (2015) was divided into two equal parts along the anterior-posterior axis, creating aSTS and pSTS parcels based on prior research showing distinct anatomical clusters in response to social interactions in posterior and anterior STS (McMahon et al., 2023). **The EVC ROI was created by combining the v1d, v1v, v2d, v2v, v3d, and v3v parcels from Wang et al., (2015).

| ROI Name | Parcel Source | Functional Localizer Task | ROI definition |
| --- | --- | --- | --- |
| pSTS (L and R) | (Deen et al., 2015)* | SI-pSTS localizer | Top 10% voxels in contrast: ‘interaction’ - ‘no-interaction’ |
| aSTS (L and R) | (Deen et al., 2015)* | SI-pSTS localizer | Top 10% voxels in contrast: ‘interaction’ - ‘no-interaction’ |
| TPJ (L and R) | (Dufour et al., 2013) | ToM localizer | Top 10% voxels in contrast: ‘belief’ - ‘photo’ |
| mmPFC | (Dufour et al., 2013) | ToM localizer | Top 10% voxels in contrast: ‘belief’ - ‘photo’ |
| dmPFC | (Dufour et al., 2013) | ToM localizer | Top 10% voxels in contrast: ‘belief’ - ‘photo’ |
| vmPFC | (Dufour et al., 2013) | ToM localizer | Top 10% voxels in contrast: ‘belief’ - ‘photo’ |
| MT (L and R) | (Wang et al., 2015) | SI-pSTS localizer | Top 10% voxels in ‘no-interaction’ - ‘rest’ |
| EVC (L and R) | (Wang et al., 2015)** | - | All voxels in anatomical mask |

##### ROI-wise RSA

For each region of interest (ROI), neural activity patterns were extracted for every condition (video) in the Main Task, and representational dissimilarity matrices (RDMs) were computed using 1 - Pearson correlation for each pair of videos. These ROI-specific RDMs were then compared to computational model RDMs and behavioral data RDMs using Spearman’s rank correlation. In deviation from the preregistered plan, significance testing was conducted at the group level rather than within individual participants. This approach assesses whether the mean RSA correlation (between model and neural RDMs) is consistently greater than zero across participants, providing a direct test of population-level effects. Significance was assessed using one-tailed signed permutation testing with FDR correction (α = 0.05). To test whether correlations between model and neural RDMs differed significantly between models, two-tailed signed permutation tests were conducted with FDR correction (α = 0.05) across model comparisons within each ROI. In addition to standard RSA, we also performed variance partitioning to quantify the unique variance in neural representational structure explained by each model or behavioral feature while controlling for the others. This was done using semi-partial Spearman correlations, and significance testing followed the same signed permutation and FDR correction procedures described above. In an exploratory analysis to assess whether anatomical overlap between right pSTS and right TPJ influenced model–brain correspondence, we repeated the ROI-based variance partitioning analyses after excluding voxels shared between the two ROIs.

##### Time-resolved ROI-wise RSA

In an exploratory analysis, neural responses were modeled separately for each of five consecutive 2s time windows within the 10s videos. For each window, voxel-wise beta estimates were computed and used to construct ROI-wise RDMs. Model-brain correspondence was assessed using semi-partial Spearman correlations, estimating the unique variance explained by SocialGNN or SIMPLE while controlling for the other model. For each time window, significance of model-brain correspondence relative to zero was assessed using one-tailed signed permutation testing with FDR correction, following the same procedure as the standard ROI-wise RSA analyses. To assess whether model correspondence varied across time, repeated-measures ANOVAs were conducted with factors model (SocialGNN, SIMPLE) and time window (five levels), separately for each ROI. These analyses tested for model-by-time interactions.

#### 2.3 Computational Models and Behavioral Judgements

##### Human social judgements

To compare human social evaluations on the PHASE dataset to different models, we used human judgements of the social relationship between the two agents collected in our previous study (Malik & Isik, 2023) from the Prolific (https://www.prolific.com/) online platform. Informed consent was obtained from all participants before the experiment. In each trial, the participants were asked to rate the relationship between the two creatures. Each participant rated 23 randomly ordered videos, including a random subset of 20 from the dataset and the three example videos shown in the instruction phase of the study as catch trials. In the current experiment, videos were picked from the generalization set only. For this set, human ratings were collected from 103 participants (ages: 19–72, mean age = 37; sex: 55 Female, 46 Male, 2 Unspecified). After exclusions, the study was left with 67 participants for the generalization set, with 10–19 ratings per video (median number of ratings = 10). For more details see Malik & Isik, (2023). Note that the videos presented to collect these ratings were from the full videos and the 10s versions were checked for consistency (see Stimuli section above).

We created behavior representations for each video using the counts of "friendly", "neutral", and "adversarial" ratings given to that video by all participants, and standardized this array to sum to 1. For example, if a video was rated friendly by 8 participants, adversarial by 2, and neutral by 0, then the human ratings representation for that video would be (0.8, 0, 0.2). This representation captures the ambiguity in human ratings. Using these, we again calculated 1 - Pearson correlation between the representations of each pair of videos to create the RDM.

##### Post-scan behavioral ratings

In supplementary analyses, we additionally constructed an RDM from post-scan social relationship ratings provided by the fMRI participants themselves (see Methods: Post-Scan Behavioral Ratings). For each video, participants’ categorical judgments ("friendly", "neutral", or "adversarial") were encoded as a one-hot vector. Pairwise dissimilarities between videos were then computed as 1 − Pearson correlation between these representations, yielding a post-scan ratings RDM. This RDM was used only in supplementary analyses to assess the relationship between each participant’s neural representations and their own social judgments.

##### SocialGNN

SocialGNN is a graph-neural-network based bottom-up model that incorporates relational inductive biases to recognize social interactions from visual scenes (Malik & Isik, 2023). For each video, it takes in a graph representation of the visual information for each frame. The nodes in these input graphs are the visual properties of the entities (agents and objects, including 2D position, 2D velocity, angle, size, and whether that entity is an agent or object) in the frame and the edges represent physical contact between the entities. Each graph also gets contextual information in the form of walls and landmarks coordinates. For each video, this sequence of graphs acts as the input to SocialGNN. Importantly, all input features and edges could be extracted from purely visual input (though here we take advantage of the annotations provided with the dataset). The overall network architecture is similar to a recurrent neural network (RNN) that at each time step processes new input graphs (GNN) and combines it with the learned representations from prior timesteps (LSTM) (Fig. 2c). The model combines these representations across time and uses the representation at the final time step to predict the type of social interaction as "friendly", "neutral", or "adversarial" via a linear classifier. See Malik & Isik, (2023) for more details. The model was trained with the following parameters: 100 epochs, batch size of 20, learning rate of 1e-3, L2 regularization parameter of 0.05, LSTM layer size of 16, and GNN edge update function using a linear layer of size 64. Here, SocialGNN is trained on 400 videos (each trimmed to 10 seconds, similar to our 50 selected videos), and predictions are made on the 50 selected videos. To construct the model’s RDM, we used the array representation from the final RNN step for each video and computed 1 - Pearson correlation between all video pairs. In exploratory analyses, we additionally constructed an RDM using the model’s final classifier output probabilities for the three interaction categories, instead of the embeddings from the final RNN step, and analyzed these using the same RSA and variance-partitioning procedures described above.

##### SIMPLE

This is a generative inverse-planning model (Fig. 2d) (Netanyahu et al., 2021). Briefly, for each video, SIMPLE generates hypotheses about the possible physical and social goals and relationships of the agents, simulates trajectories corresponding to these hypotheses, and compares the simulated trajectories to the observed trajectories of the entities in the video. The model selects the relationship with the best match between simulated and observed trajectories. To reduce its search space, SIMPLE first shortlists a set of hypotheses using bottom-up cues. These hypotheses are then updated over multiple iterations of trajectory simulation. For trajectory simulation, SIMPLE uses a hierarchical planner for each agent that, for each hypothesis, generates subgoals and actions based on the agent’s own mental state and partial view of the environment. These actions are then executed via a physics engine to produce the simulated trajectory. For more details refer to the model paper (Netanyahu et al., 2021). To construct this model’s RDMs, we used the predicted probabilities for each relationship category as the representation for each of the 50 selected videos. The RDM was then computed using 1 - Pearson correlation between the representations of all video pairs. In exploratory analyses, we tested alternative representations for SIMPLE beyond the social relationship probability vector used in the main analyses. These included a combined representation of predicted social relationship and physical goal probabilities, as well as distances between observed trajectories and the model’s internally simulated trajectories for shortlisted hypotheses; all representations were analyzed using the same RSA and variance-partitioning procedures described above.

##### ControlRNN

ControlRNN has the same broad RNN architecture and input information as SocialGNN but lacks the graph structure and graph processing (Malik & Isik, 2023). Essentially, the node features used in the visual graphs for SocialGNN input, are instead concatenated for all entities in the scene and directly input to the LSTM. Like SocialGNN, we make predictions using this model on the 50 held-out videos used in the fMRI experiment and then take the outputs from the last LSTM step to create the model’s RDM (Fig. S4). The model was trained with the following parameters: 100 epochs, batch size of 20, learning rate of 1e-3, L2 regularization parameter of 0.01, LSTM layer size of 16, and input feature size 52.

##### Motion Energy

As a low-level visual control, we compared neural responses to the output of a motion energy model (Adelson & Bergen, 1985) (Fig. 2b). Motion energy features were computed for each frame in each video from the *pymoten* package (Nunez-Elizalde et al., 2024). In deviation from the preregistered plan, we first applied principal component analysis (PCA) to these framewise features (instead of using the full 1840 dimensions) to reduce redundancy and noise inherent in the raw motion energy features. The top 128 components, explaining approximately 90% of the variance, were retained and then averaged across frames to obtain the motion energy representation for each video.

#### 2.4 Reliability and Noise Ceiling

Split-half reliability was calculated for the Main task for each participant. We divided repeated trials into two subsets (e.g., even and odd repetitions), computed the average response for each subset, and then calculated the correlation between these two averages for each voxel. A liberal split-half reliability measure was used as a mask on the neural data, ensuring that RSA was performed only in reliable voxels. For ROI-based analyses, voxels in a subject’s ROI were included if the subject-specific reliability was greater than 0; for whole-brain searchlight analyses, voxels were included if mean reliability across participants was greater than 0. For ROI-based analysis, we also calculated split-half RSA to estimate the reliability of the ROI-level representational geometry in the Main task. Here, for each ROI, we split the data into even and odd repetitions, applied the split-half reliability mask to each subset, generated RDMs, and performed RSA between the splits using Spearman’s rank correlation. These correlations were Spearman–Brown corrected (Brown, 1910; Spearman, 1910). The Spearman-Brown-corrected split-half RSA correlation, averaged across participants, was used as a reference reliability estimate for ROI-level model comparisons (Fig. S1b). RSA results for ROIs where this reference reliability was not significantly above 0 are not reported.

#### 2.5 Data and Code Availability

All behavioral and neuroimaging data, model implementations, and analysis code will be made publicly available upon publication via GitHub (https://github.com/Isik-lab/SI-Neural-Computations) and the Open Science Framework (OSF).

## Supporting information

Supplemental Section

## Acknowledgments

This work was supported by NIMH R01MH132826 awarded to L.I. We would like to thank Wenshuo Qin, Elizabeth Jiwon Im, Emalie McMahon, and the F.M. Kirby Research Center staff for help with fMRI data collection, and Hannah Small for feedback on code. We would also like to thank Josh Tenenbaum, Barbara Landau, and Sam Maione for helpful discussions of this work.

## REFERENCES

Adelson, E. H., & Bergen, J. R. (1985). Spatiotemporal energy models for the perception of motion. Journal of the Optical Society of America. A, Optics and Image Science, 2(2), 284. 10.1364/JOSAA.2.000284

Baillargeon, R., Scott, R. M., & Bian, L. (2016). Psychological Reasoning in Infancy. Annual Review of Psychology, 67, 159–186. 10.1146/ANNUREV-PSYCH-010213-115033

Baker, C. L., Jara-Ettinger, J., Saxe, R., & Tenenbaum, J. B. (2017). Rational quantitative attribution of beliefs, desires and percepts in human mentalizing. Nature Human Behaviour 2017 1:4, 1(4), 0064-. 10.1038/s41562-017-0064

Baron-Cohen, S., Wheelwright, S., Skinner, R., Martin, J., & Clubley, E. (2001). The Autism-Spectrum Quotient (AQ): Evidence from Asperger Syndrome/High-Functioning Autism, Males and Females, Scientists and Mathematicians. Journal of Autism and Developmental Disorders, 31(1), 5–17. 10.1023/A:1005653411471

Battaglia, P. W., Hamrick, J. B., & Tenenbaum, J. B. (2013). Simulation as an engine of physical scene understanding. Proceedings of the National Academy of Sciences of the United States of America, 110(45), 18327–18332. 10.1073/pnas.1306572110

Bear, D. M., Fan, C., Mrowca, D., Li, Y., Alter, S., Nayebi, A., Schwartz, J., Fei-Fei, L., Wu, J., Tenenbaum, J. B., & Yamins, D. L. K. (2020). Learning Physical Graph Representations from Visual Scenes. http://arxiv.org/abs/2006.12373

Bi, W., Shah, A. D., Wong, K. W., Scholl, B. J., & Yildirim, I. (2025). Computational models reveal that intuitive physics underlies visual processing of soft objects. Nature Communications 2025 16:1, 16(1), 6303-. 10.1038/s41467-025-61458-x

Brown, W. (1910). Some experimental results in the correlation of mental abilities. British Journal of Psychology, 1904-1920, 3(3), 296–322. 10.1111/J.2044-8295.1910.TB00207.X;REQUESTEDJOURNAL:JOURNAL:20448295B;WGROUP:STRING:PUBLICATION

Deen, B., Koldewyn, K., Kanwisher, N., & Saxe, R. (2015). Functional Organization of Social Perception and Cognition in the Superior Temporal Sulcus. Cerebral Cortex, 25(11), 4596–4609. 10.1093/CERCOR/BHV111

Dodell-Feder, D., Koster-Hale, J., Bedny, M., & Saxe, R. (2011). fMRI item analysis in a theory of mind task. NeuroImage, 55(2), 705–712. 10.1016/J.NEUROIMAGE.2010.12.040

Dufour, N., Redcay, E., Young, L., Mavros, P. L., & Moran, J. M. (2013). Similar Brain Activation during False Belief Tasks in a Large Sample of Adults with and without Autism. PLoS ONE, 8(9), 75468. 10.1371/journal.pone.0075468

Esteban, O., Markiewicz, C. J., Blair, R. W., Moodie, C. A., Isik, A. I., Erramuzpe, A., Kent, J. D., Goncalves, M., DuPre, E., Snyder, M., Oya, H., Ghosh, S. S., Wright, J., Durnez, J., Poldrack, R. A., & Gorgolewski, K. J. (2018). fMRIPrep: a robust preprocessing pipeline for functional MRI. Nature Methods 2018 16:1, 16(1), 111–116. 10.1038/s41592-018-0235-4

Fischer, J., Mikhael, J. G., Tenenbaum, J. B., & Kanwisher, N. (2016). Functional neuroanatomy of intuitive physical inference. Proceedings of the National Academy of Sciences of the United States of America, 113(34). 10.1073/pnas.1610344113

Gandhi, K., Stojnic, G., Lake, B. M., & Dillon, M. R. (2021). Baby Intuitions Benchmark (BIB): Discerning the goals, preferences, and actions of others. Advances in Neural Information Processing Systems, 34, 9963–9976. https://www.youtube.com/watch?v=VTNmLt7QX8E

Gao, T., Scholl, B. J., & McCarthy, G. (2012). Dissociating the Detection of Intentionality from Animacy in the Right Posterior Superior Temporal Sulcus. Journal of Neuroscience, 32(41), 14276–14280. 10.1523/JNEUROSCI.0562-12.2012

Geraci, A., Surian, L., Tina, L. G., & Hamlin, J. K. (2025). Human newborns spontaneously attend to prosocial interactions. Nature Communications 2025 16:1, 16(1), 6304-. 10.1038/s41467-025-61517-3

Gorgolewski, K., Burns, C. D., Madison, C., Clark, D., Halchenko, Y. O., Waskom, M. L., & Ghosh, S. S. (2011). Nipype: A flexible, lightweight and extensible neuroimaging data processing framework in Python. Frontiers in Neuroinformatics, 5, 12318. 10.3389/FNINF.2011.00013/ABSTRACT

Gulban, O. F., Nielson, D., lee, john, Poldrack, R., Gorgolewski, C., Vanessasaurus, & Markiewicz, C. (2022). poldracklab/pydeface: PyDeface v2.0.2. Zenodo. 10.5281/zenodo.6856482

Hafri, A., & Firestone, C. (2021). The perception of relations. Trends in Cognitive Sciences, 25(6), 475–492.

Heider, F., & Simmel, M. (1944). An experimental study of apparent behavior. The American Journal of Psychology, 57(2), 243–259.

Ho, M. K., Saxe, R., & Cushman, F. (2022). Planning with Theory of Mind. Trends in Cognitive Sciences, 26(11), 959–971. 10.1016/j.tics.2022.08.003

Isik, L., Koldewyn, K., Beeler, D., & Kanwisher, N. (2017). Perceiving social interactions in the posterior superior temporal sulcus. Proceedings of the National Academy of Sciences of the United States of America, 114(43), E9145–E9152. 10.1073/pnas.1714471114

Jamali, M., Grannan, B. L., Fedorenko, E., Saxe, R., Báez-Mendoza, R., & Williams, Z. M. (2021). Single-neuronal predictions of others’ beliefs in humans. Nature 2021 591:7851, 591(7851), 610–614. 10.1038/s41586-021-03184-0

Jara-Ettinger, J. (2019). Theory of mind as inverse reinforcement learning. Current Opinion in Behavioral Sciences, 29, 105–110. 10.1016/j.cobeha.2019.04.010

Jha, K., Le, T. A., Jin, C., Kuo, Y. L., Tenenbaum, J. B., & Shu, T. (2024). Neural Amortized Inference for Nested Multi-Agent Reasoning. Proceedings of the AAAI Conference on Artificial Intelligence, 38(1), 530–537. 10.1609/aaai.v38i1.27808

Jin, C., Wu, Y., Cao, J., Xiang, J., Kuo, Y.-L., Hu, Z., Ullman, T., Torralba, A., Tenenbaum, J. B., & Shu, T. (2024). MMToM-QA: Multimodal Theory of Mind Question Answering. Proceedings of the Annual Meeting of the Association for Computational Linguistics, 1, 16077–16102. http://arxiv.org/abs/2401.08743

Kay, K. N., Rokem, A., Winawer, J., Dougherty, R. F., & Wandell, B. A. (2013). GLMdenoise: A fast, automated technique for denoising task-based fMRI data. Frontiers in Neuroscience, (7 DEC). 10.3389/FNINS.2013.00247

Kim, H., Sclar, M., Zhi-Xuan, T., Ying, L., Levine, S., Liu, Y., Tenenbaum, J. B., & Choi, Y. (2025). Hypothesis-Driven Theory-of-Mind Reasoning for Large Language Models. http://arxiv.org/abs/2502.11881

K.J. Gorgolewski, O. Esteban, C.J. Markiewicz, E. Ziegler, D.G. Ellis, M.P. Notter, D. Jarecka, H. Johnson, C. Burns, & A. Manhães-Savio, et al. (2018). nipy/nipype: 1.8.3. 10.5281/ZENODO.6834519

Koster-Hale, J., Richardson, H., Velez, N., Asaba, M., Young, L., & Saxe, R. (2017). Mentalizing regions represent distributed, continuous, and abstract dimensions of others’ beliefs. NeuroImage, 161, 9–18. 10.1016/J.NEUROIMAGE.2017.08.026

Kriegeskorte, N., Mur, M., & Bandettini, P. (2008). Representational similarity analysis - connecting the branches of systems neuroscience. Frontiers in Systems Neuroscience, 2(NOV), 249. 10.3389/NEURO.06.004.2008/BIBTEX

Krupenye, C., & Hare, B. (2018). Bonobos prefer individuals that hinder others over those that help. Current Biology, 28(2), 280–286.

Liu, S., Karakose-Akbiyik, S., Outa, J., & Kim, M. J. (2025). How physical information is used to make sense of the psychological world. Nature Reviews Psychology 2025 5:1, 5(1), 59–73. 10.1038/s44159-025-00514-1

Malik, M., & Isik, L. (2023). Relational visual representations underlie human social interaction recognition. Nature Communications 2023 14:1, 14(1), 1–11. 10.1038/s41467-023-43156-8

Malik, M., Kim, M., Liu, S., Shu, T., & Isik, L. (2025). Neural representations and computations underlying human social evaluations from visual stimuli. 10.17605/OSF.IO/B6E2Q

McMahon, E., Bonner, M. F., & Isik, L. (2023). Hierarchical organization of social action features along the lateral visual pathway. Current Biology, 33(23), 5035–5047.e8. 10.1016/J.CUB.2023.10.015

McMahon, E., & Isik, L. (2023). Seeing social interactions. Trends in Cognitive Sciences, 0(0). 10.1016/J.TICS.2023.09.001

Netanyahu, A., Shu, T., Katz, B., Barbu, A., & Tenenbaum, J. B. (2021). PHASE: PHysically-grounded Abstract Social Events for Machine Social Perception. Proceedings of the AAAI Conference on Artificial Intelligence, 35(1), 845–853. 10.1609/AAAI.V35I1.16167

Nunez-Elizalde, A. O., Deniz, F., la Tour, T., di Oleggio Castello, M., & Gallant, J. L. (2024). pymoten: scientific python package for computing motion energy features from video. Zenodo. 10.5281/zenodo.13328756

Papeo, L. (2020). Twos in human visual perception. Cortex, 132, 473–478. 10.1016/J.CORTEX.2020.06.005

Piloto, L. S., Weinstein, A., Battaglia, P., & Botvinick, M. (2022). Intuitive physics learning in a deep-learning model inspired by developmental psychology. Nature Human Behaviour 2022 6:9, 6(9), 1257–1267. 10.1038/s41562-022-01394-8

Powell, L. J., & Spelke, E. S. (2018). Human infants’ understanding of social imitation: Inferences of affiliation from third party observations. Cognition, 170, 31–48. 10.1016/j.cognition.2017.09.007

Pramod, R. T., Mieczkowski, E., Fang, C. X., Tenenbaum, J. B., & Kanwisher, N. (2025). Decoding predicted future states from the brain’s “physics engine”. Science Advances, 11(22), eadr7429–eadr7429. 10.1126/sciadv.adr7429

Prince, J. S., Charest, I., Kurzawski, J. W., Pyles, J. A., Tarr, M. J., & Kay, K. N. (2022). Improving the accuracy of single-trial fMRI response estimates using GLMsingle. ELife, 11. 10.7554/ELIFE.77599

Qin, W., Malik, M., & Isik, L. (2025). Relational Information Predicts Human Behavior and Neural Responses to Complex Social Scenes. Proceedings of the Annual Meeting of the Cognitive Science Society, 47. https://escholarship.org/uc/item/4680v4ws

Saxe, R., & Kanwisher, N. (2003). People thinking about thinking people: The role of the temporo-parietal junction in “theory of mind.” NeuroImage, 19(4), 1835–1842. 10.1016/S1053-8119(03)00230-1

Shi, H., Ye, S., Fang, X., Jin, C., Isik, L., Kuo, Y.-L., & Shu, T. (2025). MuMA-ToM: Multi-modal Multi-Agent Theory of Mind. Proceedings of the AAAI Conference on Artificial Intelligence, 39(2), 1510–1519. http://arxiv.org/abs/2408.12574

Shu, T., Bhandwaldar, A., Gan, C., Smith, K., Liu, S., Gutfreund, D., Spelke, E., Tenenbaum, J., & Ullman, T. (2021). Agent: A benchmark for core psychological reasoning. International Conference on Machine Learning, 9614–9625.

Skerry, A. E., & Saxe, R. (2015). Neural Representations of Emotion Are Organized around Abstract Event Features. Current Biology, 25(15), 1945–1954. 10.1016/j.cub.2015.06.009

Spearman, C. (1910). Correlation calculated from faulty data. British Journal of Psychology, 1904-1920, 3(3), 271–295. 10.1111/J.2044-8295.1910.TB00206.X

Spelke, E. S. (2022). Agents. What Babies Know, 248–299. 10.1093/OSO/9780190618247.003.0007

Ullman, T. D., Baker, C. L., Macindoe, O., Evans, O., Goodman, N. D., & Tenenbaum, J. B. (2009). Help or Hinder: Bayesian Models of Social Goal Inference. Advances in Neural Information Processing Systems, 22.

Varrier, R. S., Su, Z., Liang, Q., Benson, T. G., Jolly, E., & Finn, E. S. (2025). Shared and individual tuning curves for social perception. BioRxiv, 2025.01.19.633772. 10.1101/2025.01.19.633772

Walbrin, J., Downing, P., & Koldewyn, K. (2018). Neural responses to visually observed social interactions. Neuropsychologia, 112, 31–39. 10.1016/J.NEUROPSYCHOLOGIA.2018.02.023

Wang, L., Mruczek, R. E. B., Arcaro, M. J., & Kastner, S. (2015). Probabilistic Maps of Visual Topography in Human Cortex. Cerebral Cortex, 25(10), 3911–3931. 10.1093/CERCOR/BHU277

Woo, B. M., Tan, E., & Hamlin, J. K. (2022). Human Morality Is Based on an Early-Emerging Moral Core. Https://Doi.Org/10.1146/Annurev-Devpsych-121020-023312, 4(1). 10.1146/ANNUREV-DEVPSYCH-121020-023312

Yildirim, I., Belledonne, M., Freiwald, W., & Tenenbaum, J. (2020). Efficient inverse graphics in biological face processing. Science Advances, 6(10). 10.1126/sciadv.aax5979

Zhang, Z., Jin, C., Jia, M. Y., Zhang, S., & Shu, T. (2026). AutoToM: Scaling Model-based Mental Inference via Automated Agent Modeling. NeurIPS, 2025. http://arxiv.org/abs/2502.15676

Zhou, M., Gong, Z., Dai, Y., Wen, Y., Liu, Y., & Zhen, Z. (2023). A large-scale fMRI dataset for human action recognition. Scientific Data, 10(1). 10.1038/s41597-023-02325-6

