## Supplemental Section for "Bottom-up and generative computations uniquely explain neural responses across the social brain"

### SUPPLEMENTARY SECTION

#### S1. ROI-RSA results across all ROIs

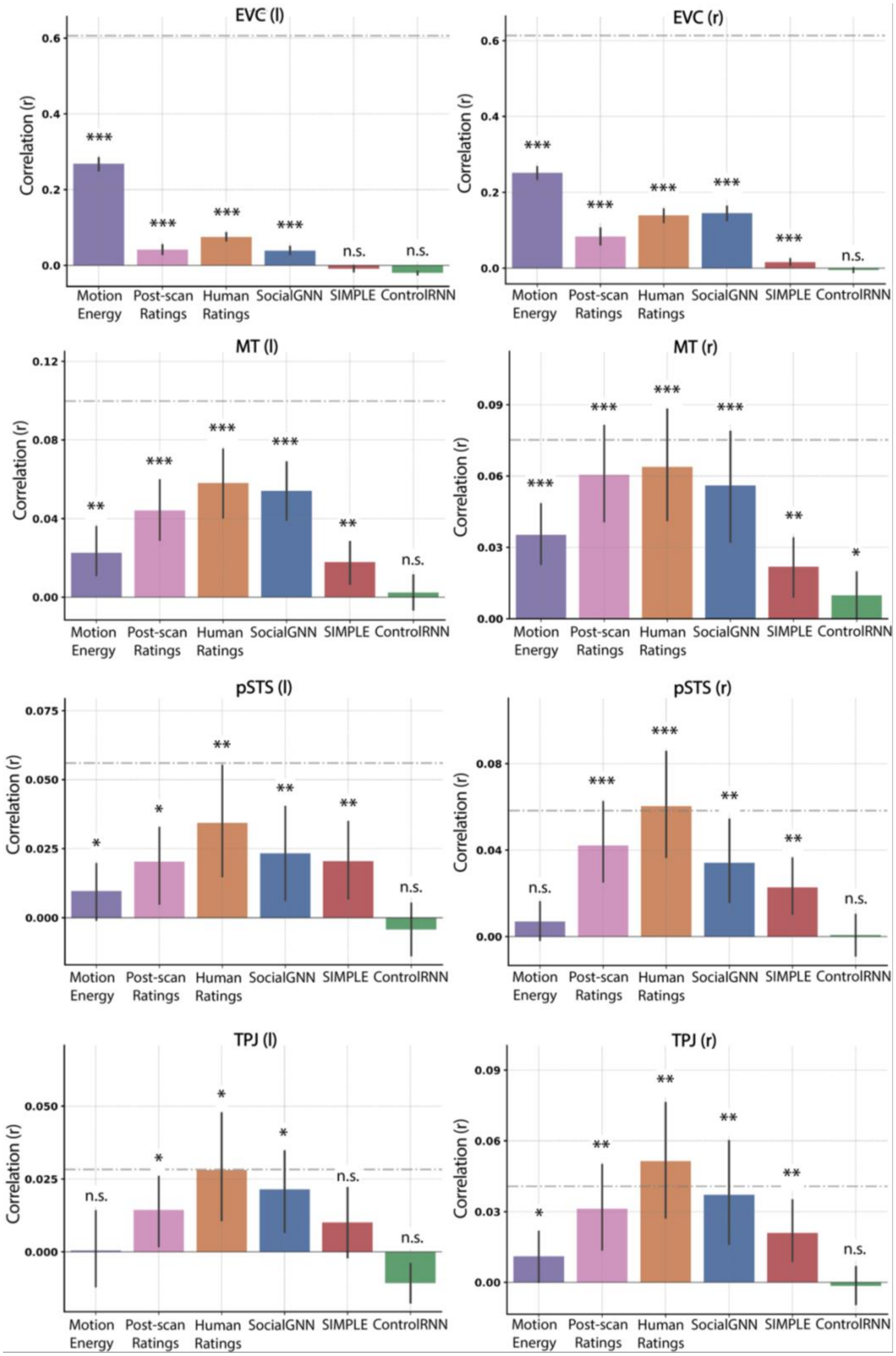


***Fig. S1a.*** *ROI-based representational similarity analysis results for all ROIs. Bar height indicates the mean Spearman correlation (across participants) between neural RDMs and model RDMs,*

*including motion energy, human ratings (from independent raters and from fMRI participants), SocialGNN, SIMPLE, and a non-relational control model. Error bars represent 95% confidence intervals. Asterisks denote significance levels (p < 0.001 ***, p < 0.01 **, p < 0.05 *); n.s., not significant, evaluated using one-tailed signed permutation testing with FDR correction across models within each ROI (α = 0.05). The gray dashed line indicates the mean split-half RSA reliability for each ROI (spearman-brown corrected). ROIs with reliability not significantly above zero are indicated.*

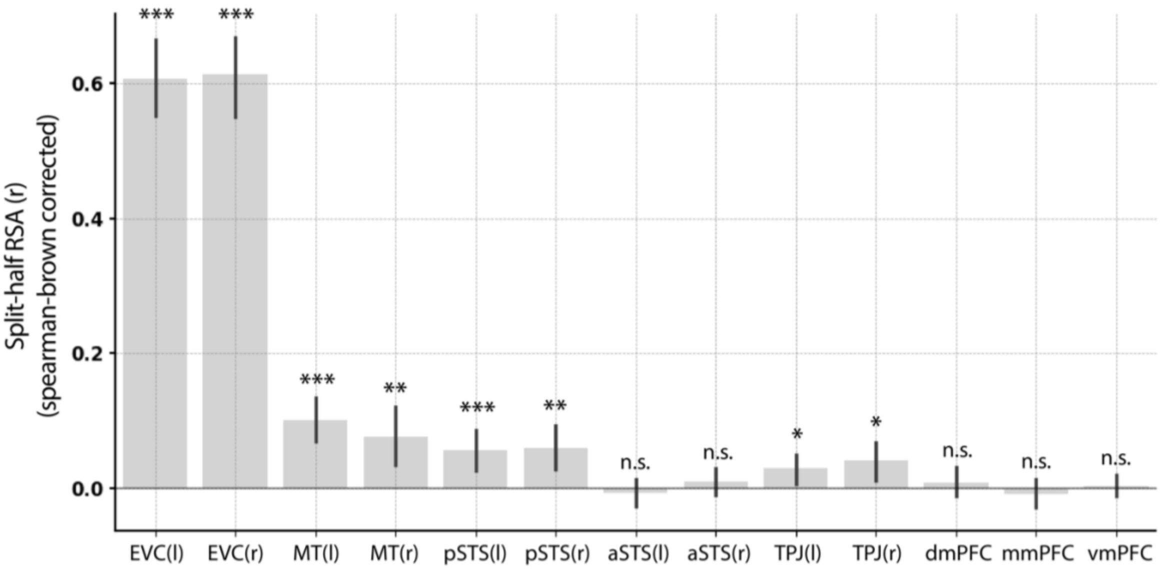


***Fig. S1b.*** *Split-half RSA-based reliability estimates for each ROI, computed using only reliable voxels. Bars show the mean split-half correlation across participants after Spearman–Brown correction; error bars indicate 95% confidence intervals across participants.*

***Table S1.*** *ROI-wise p-values testing whether correlations between model and neural (ROI) RDMs differ significantly between models. Two-tailed signed permutation tests were used, with FDR correction across model comparisons within each ROI. Cell backgrounds are shaded according to: p < 0.001 black, p < 0.01 dark gray, p < 0.05 light gray, n.s. no color.*

| | Human ratings vs SIMPLE | Human ratings vs SocialGNN | Human ratings vs ControlRNN | Human ratings vs Motion energy | Motion energy vs SIMPLE | Motion energy vs SocialGNN | Motion energy vs ControlRNN | SIMPLE vs ControlRNN | SocialGNN vs SIMPLE | SocialGNN vs ControlRNN |
|---|---|---|---|---|---|---|---|---|---|---|
| **EVC (l)** | 0.0002 | 0.0002 | 0.0002 | 0.0002 | 0.0002 | 0.0002 | 0.0002 | 0.011 | 0.0002 | 0.0002 |
| **EVC (r)** | 0.0002 | 0.1952 | 0.0002 | 0.0002 | 0.0002 | 0.0002 | 0.0002 | 0.0002 | 0.0002 | 0.0002 |
| **MT (l)** | 0.0005 | 0.4903 | 0.0005 | 0.0028 | 0.5377 | 0.0037 | 0.0094 | 0.0877 | 0.0005 | 0.0005 |
| **MT (r)** | 0.002 | 0.3231 | 0.002 | 0.0996 | 0.1554 | 0.156 | 0.0144 | 0.156 | 0.011 | 0.002 |
| **pSTS (l)** | 0.1296 | 0.2659 | 0.047 | 0.1296 | 0.3197 | 0.2802 | 0.0735 | 0.047 | 0.7574 | 0.0735 |
| **pSTS (r)** | 0.0025 | 0.0103 | 0.002 | 0.002 | 0.088 | 0.0163 | 0.2911 | 0.0048 | 0.182 | 0.0025 |
| **TPJ (l)** | 0.0435 | 0.3935 | 0.004 | 0.0836 | 0.3935 | 0.08 | 0.1543 | 0.0293 | 0.2012 | 0.004 |
| **TPJ (r)** | 0.0105 | 0.0851 | 0.008 | 0.0172 | 0.2314 | 0.0817 | 0.087 | 0.0105 | 0.1064 | 0.0105 |

#### S2. Variance uniquely explained by SocialGNN, SIMPLE, and Human social ratings after controlling for Motion energy

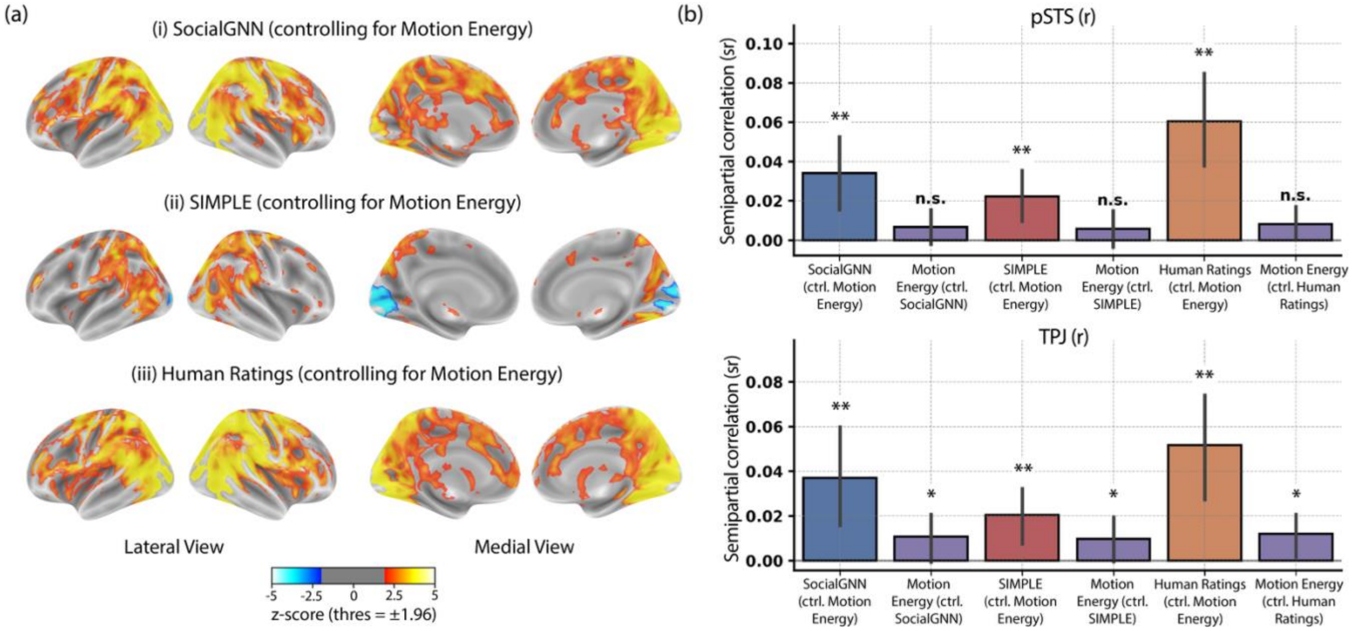


***Fig. S2:*** *(A) Whole-brain results (group-level z-scored maps), computed using semi-partial RSA between neural RDMs and SocialGNN, SIMPLE, or human ratings while controlling for motion energy. Statistical significance was determined using two-tailed signed permutation testing with FDR correction across voxels (α = 0.05). Maps are shown on inflated cortical surfaces (medial and lateral views). (B) ROI-based results in pSTS (r) and TPJ (r). Bar height indicates the mean semi-partial correlation across participants, with error bars representing 95% confidence intervals. Asterisks denote significance levels (p < 0.001 ***, p < 0.01 **, p < 0.05 *); n.s., not significant. Significance was evaluated using one-tailed signed permutation testing with FDR correction (α = 0.05).*

#### S3. Variance uniquely explained by SocialGNN, SIMPLE, and Human social ratings after controlling for early visual cortex (EVC) responses

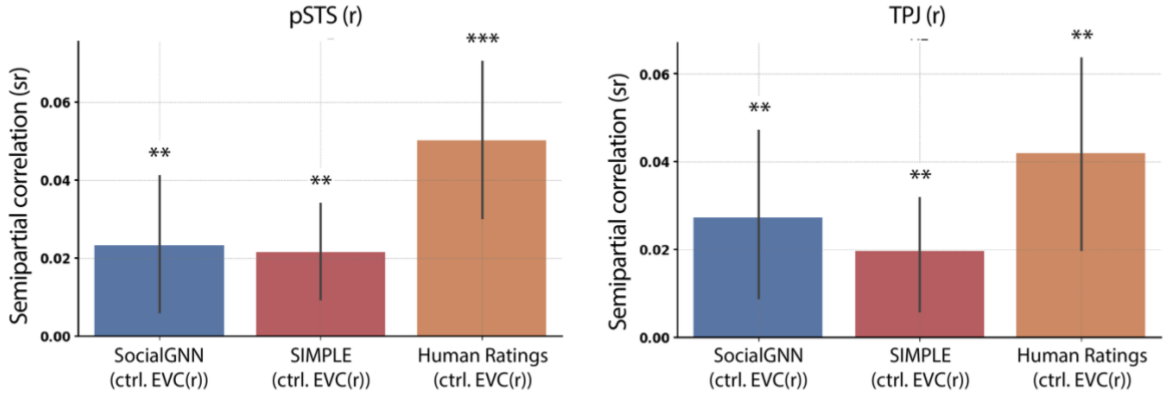


***Fig. S3:*** *Semipartial correlation results in pSTS (r) (left) and TPJ (r) (right), with early visual cortex (EVC) representations partialed out. Bar height indicates the mean semipartial Spearman*

*correlation across participants, with error bars representing 95% confidence intervals. Significance testing and annotation follow the same conventions as Fig. S2.*

#### S4. ControlRNN model architecture

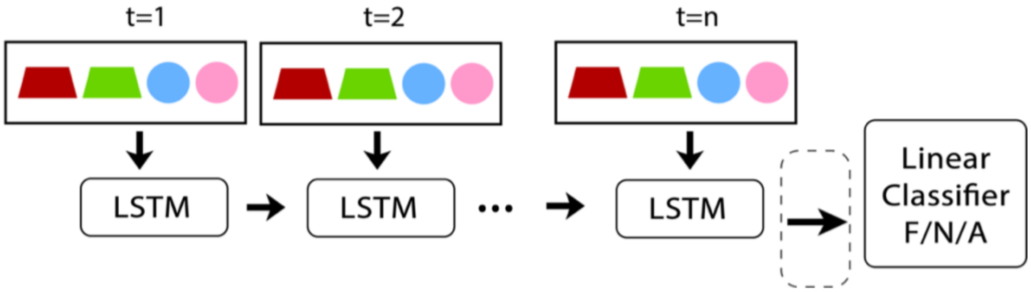


***Fig. S4: ControlRNN Model Architecture and Representation.*** *ControlRNN has the same broad RNN architecture and input information as SocialGNN but lacks the graph structure and graph processing. Essentially, the node features used in the visual graphs for SocialGNN input, are instead concatenated for all entities in the scene and directly input to the LSTM. Like SocialGNN, the final LSTM hidden state is passed to a linear classifier to predict the relationship category ("friendly", "neutral", or "adversarial") and is used as the ControlRNN representation for each video (circled with dotted lines).*

#### S5. Alternate representations from SocialGNN

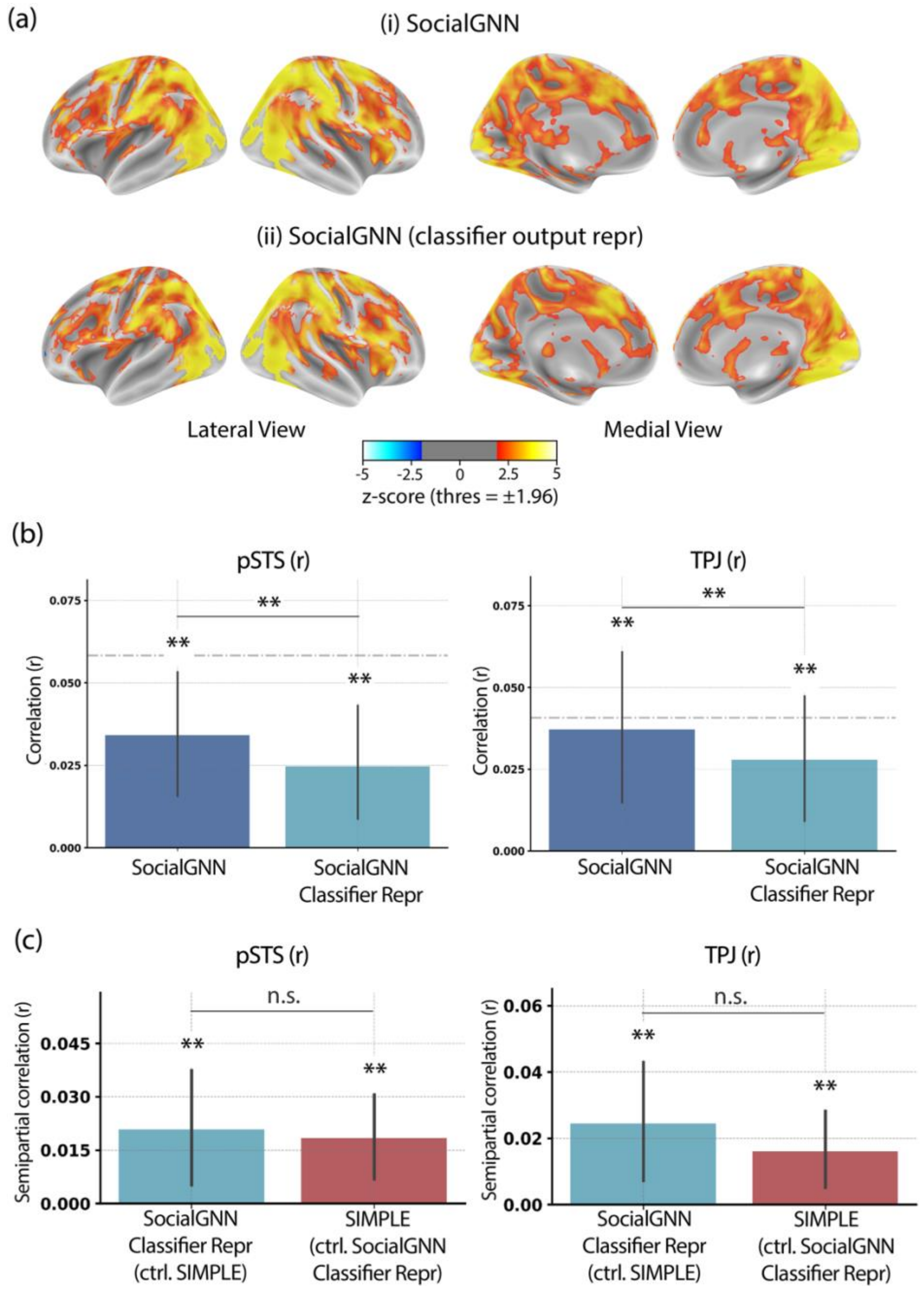


***Fig. S5:*** *(a) Whole-brain semi-partial RSA results comparing two representations derived from SocialGNN: the standard SocialGNN representation and the classifier output representation. Group-level z-scored maps were computed using RSA between neural RDMs and each representation. Results are shown on inflated cortical surfaces (medial and lateral views). Significance testing and annotation follow the same conventions as Fig. S2. (b) ROI-based RSA results in pSTS (r) and TPJ (r) comparing the standard SocialGNN representation with the classifier output representation. Bar height indicates the mean correlation across participants, with error bars representing 95% confidence intervals. Significance testing and annotation follow the same conventions as Fig. S2. (c) ROI-based unique variance (semi-partial RSA) results comparing the alternative SocialGNN representations and SIMPLE, showing unique variance explained by each model after controlling for the other. Bar height indicates the mean semi-partial correlation across*

*participants, with error bars representing 95% confidence intervals. Significance testing and annotation follow the same conventions as Fig. S2.*

#### S6. Alternate representations from SIMPLE

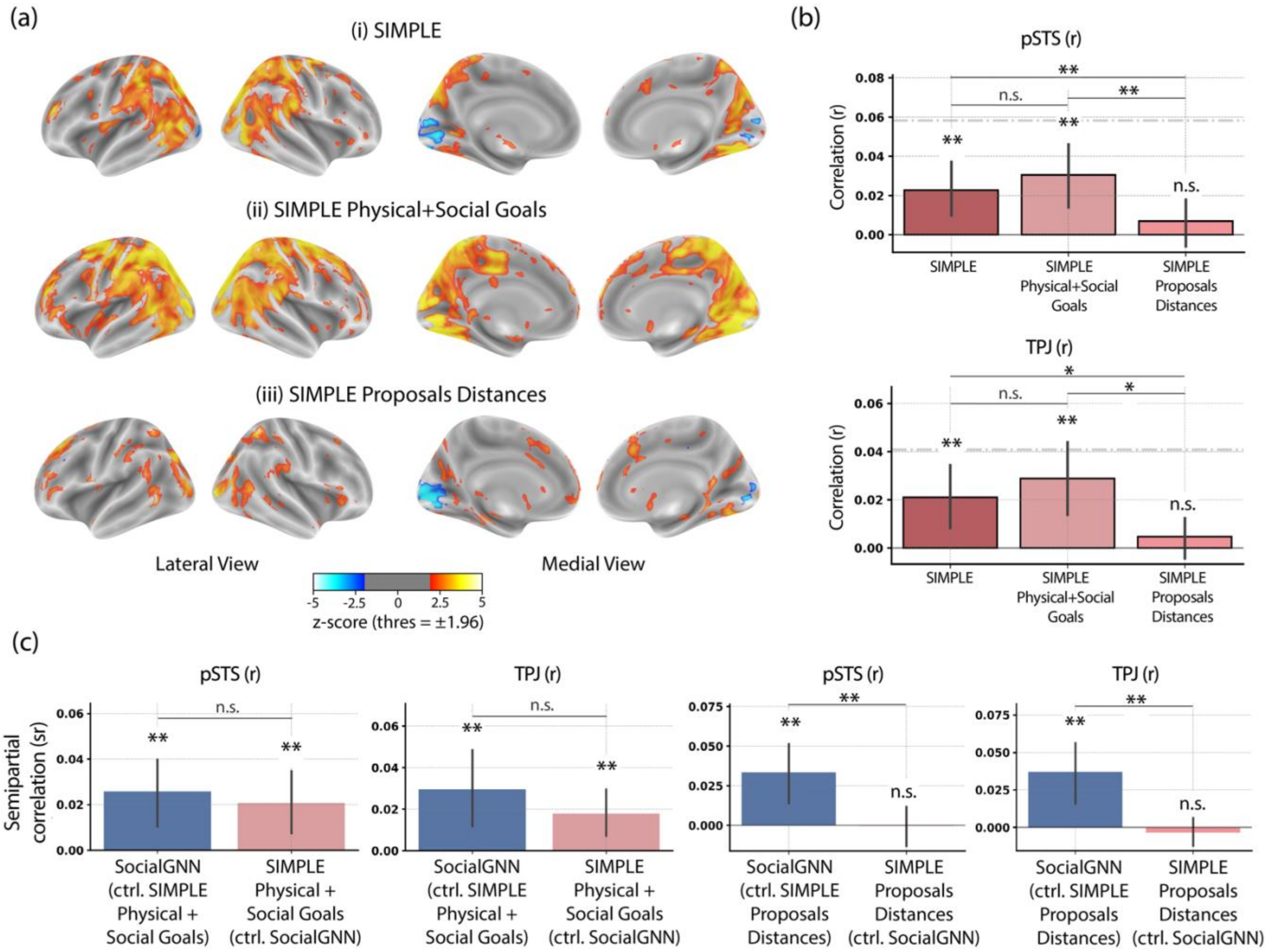


***Fig. S6.*** *(a) Whole-brain semi-partial RSA results comparing alternative representations derived from SIMPLE: the standard SIMPLE representation, a representation based on physical + social goals, and a representation based on proposal distance. Group-level z-scored maps were computed using semi-partial RSA between neural RDMs and each representation. Significance testing and annotation follow the same conventions as Fig. S2. Results are shown on inflated cortical surfaces (medial and lateral views). (b) ROI-based RSA results in pSTS (r) and TPJ (r) for the three SIMPLE representations. Bar height indicates the mean Spearman correlation across participants, with error bars representing 95% confidence intervals. Significance testing and annotation follow the same conventions as Fig. S2. (c) ROI-based unique variance (semi-partial RSA) results comparing SIMPLE representations with SocialGNN, showing the unique variance explained by each model after controlling for the other. Bar height indicates the mean semi-partial correlation across participants, with error bars representing 95% confidence intervals. Significance testing and annotation follow the same conventions as Fig. S2.*

#### S7. Effect of removing overlapping voxels between pSTS (r) and TPJ (r)

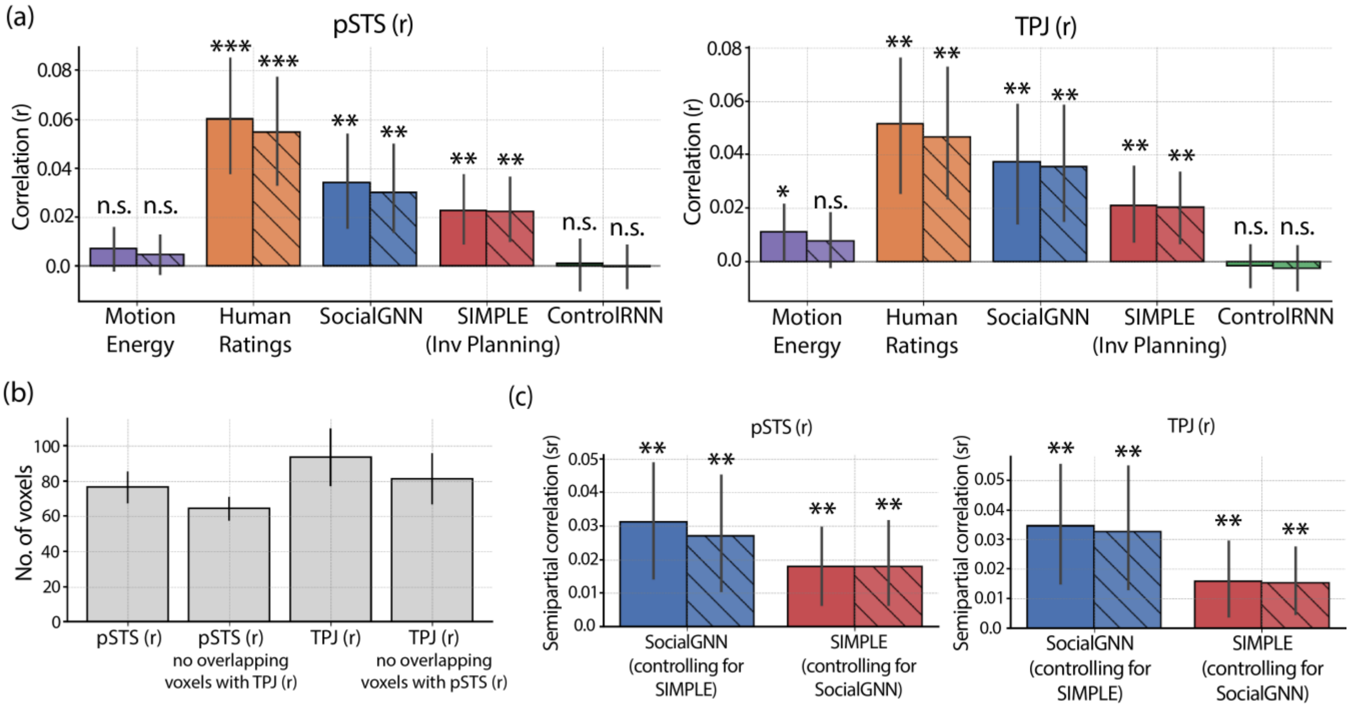


***Fig. S7:*** *(a) ROI-based RSA results in pSTS (r) and TPJ (r). Solid bars show results using the original ROIs, while hatched bars show results after excluding voxels overlapping between pSTS (r) and TPJ (r). Error bars indicate 95% confidence intervals across participants. Significance stars denote correlations significantly different from zero after FDR correction (performed separately for each ROI definition). (b) Mean number of voxels in pSTS (r) and TPJ (r) before and after removing overlapping voxels between the two regions. Error bars indicate 95% confidence intervals across participants. (c) Semi-partial RSA results in pSTS (r) and TPJ (r), showing unique variance explained by SocialGNN (controlling for SIMPLE) and SIMPLE (controlling for SocialGNN). Solid bars indicate original ROIs; hatched bars indicate ROIs with overlapping voxels removed. Error bars indicate 95% confidence intervals across participants. Significance stars denote effects surviving FDR correction, performed separately for each ROI definition.*

#### S8. ROI: supplementary details

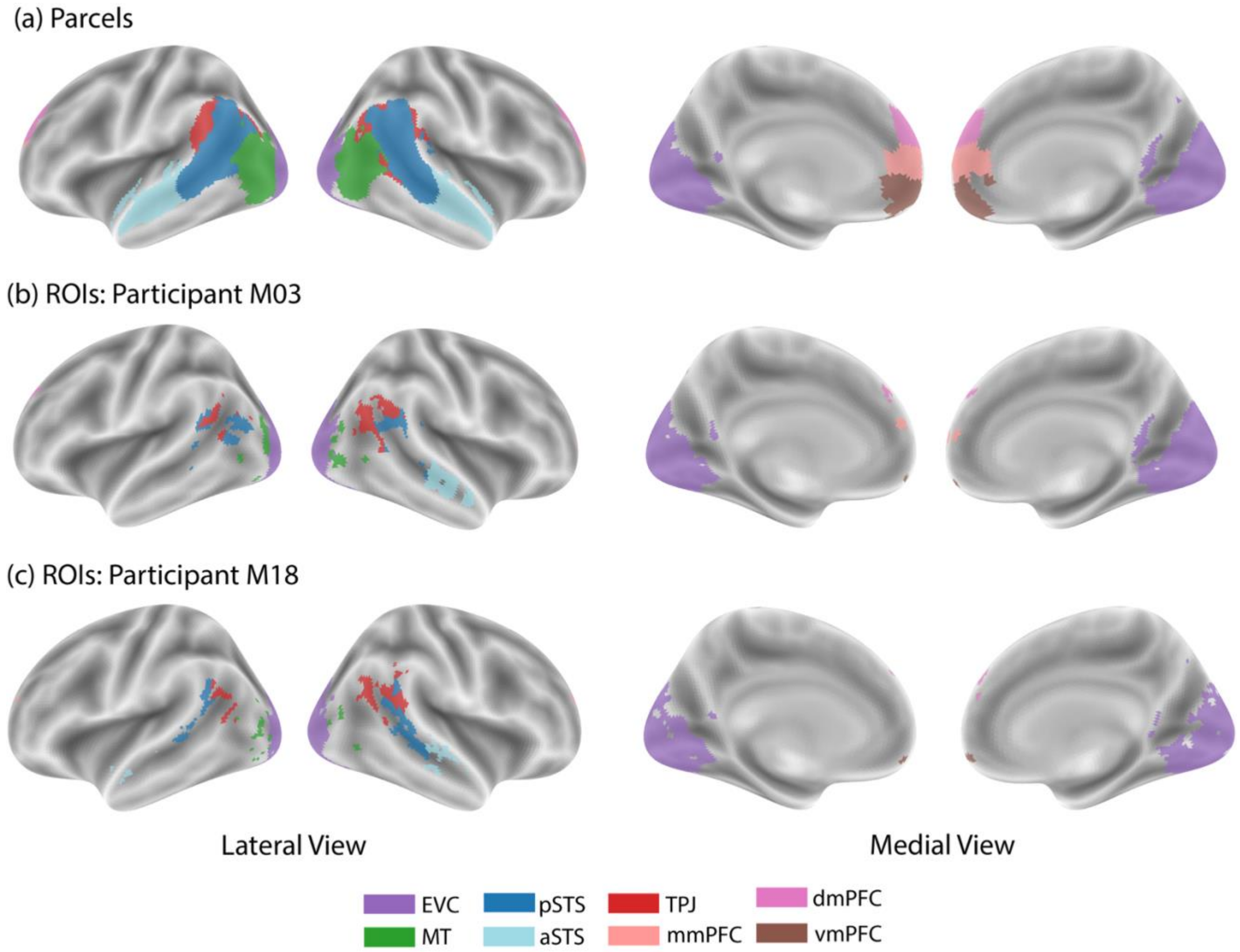


***Fig. S8. Parcels and example participant-specific ROIs***. *(a) Parcels used to define ROIs (see Table 1). (b–c) Example ROIs for two participants (M03 and M18) derived from these parcels. In areas of spatial overlap between parcels or ROIs, a single region is displayed for visualization clarity.*

***Table S2:*** *ROI sizes. Mean ± SD voxel counts for subject-specific ROIs across participants.*

| ROI | **No. of voxels** (mean ± SD) |
|---|---|
| EVC (l) | 2063.0 ± 0.0 |
| EVC (r) | 2276.0 ± 0.0 |
| MT (l) | 76.0 ± 14.6 |
| MT (r) | 33.9 ± 14.5 |
| pSTS (l) | 71.9 ± 26.1 |
| pSTS (r) | 106.1 ± 26.1 |
| aSTS (l) | 32.8 ± 15.0 |
| aSTS (r) | 64.6 ± 14.8 |
| TPJ (l) | 76.3 ± 37.8 |
| TPJ (r) | 125.7 ± 37.8 |
| dmPFC | 69.0 ± 0.0 |
| mmPFC | 53.0 ± 0.0 |
| vmPFC | 34.2 ± 4.8 |

***Table S3.*** *Mean Dice coefficients quantifying spatial overlap between ROIs across participants.*

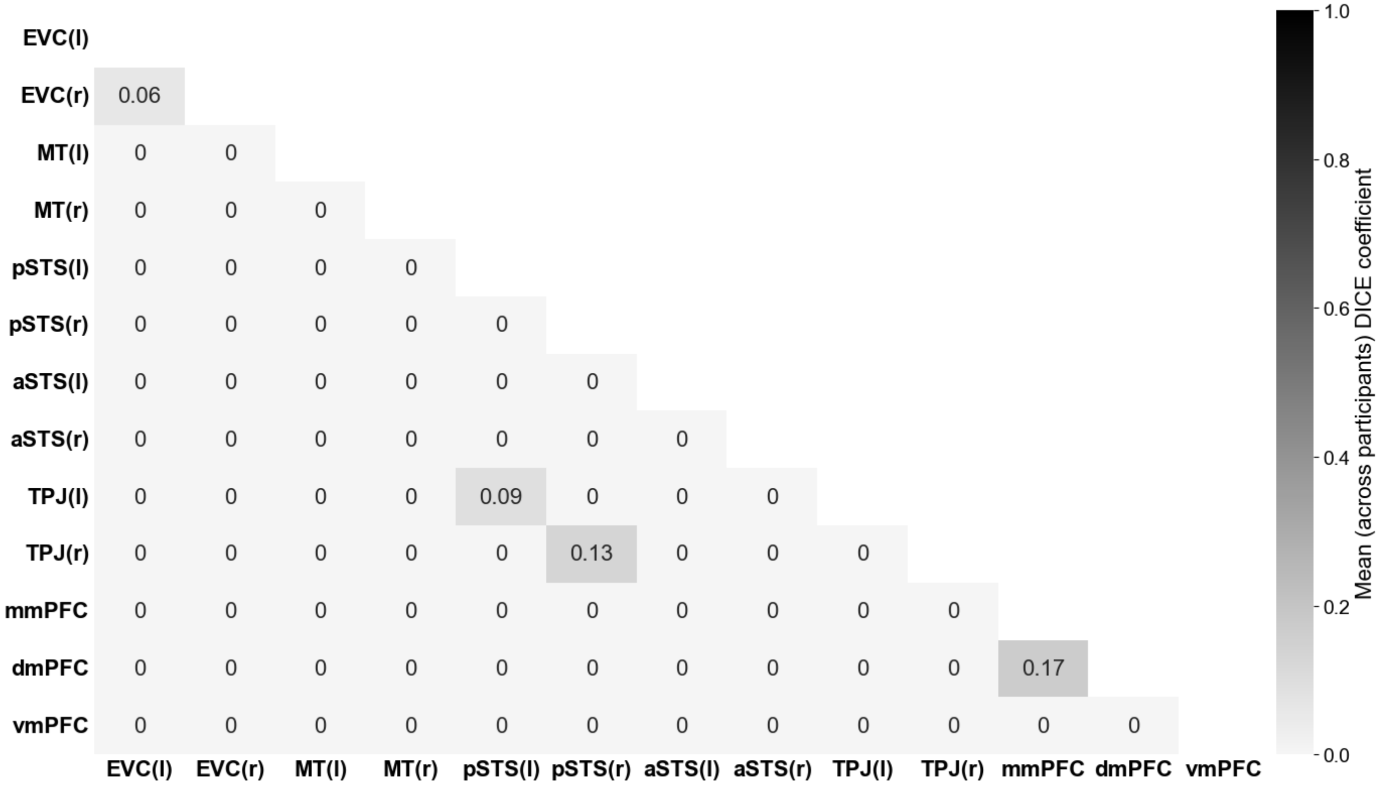


| | EVC(l) | EVC(r) | MT(l) | MT(r) | pSTS(l) | pSTS(r) | aSTS(l) | aSTS(r) | TPJ(l) | TPJ(r) | mmPFC | dmPFC | vmPFC |
|---|---|---|---|---|---|---|---|---|---|---|---|---|---|
| **EVC(l)** | | | | | | | | | | | | | |
| **EVC(r)** | 0.06 | | | | | | | | | | | | |
| **MT(l)** | 0 | 0 | | | | | | | | | | | |
| **MT(r)** | 0 | 0 | 0 | | | | | | | | | | |
| **pSTS(l)** | 0 | 0 | 0 | 0 | | | | | | | | | |
| **pSTS(r)** | 0 | 0 | 0 | 0 | 0 | | | | | | | | |
| **aSTS(l)** | 0 | 0 | 0 | 0 | 0 | 0 | | | | | | | |
| **aSTS(r)** | 0 | 0 | 0 | 0 | 0 | 0 | 0 | | | | | | |
| **TPJ(l)** | 0 | 0 | 0 | 0 | 0.09 | 0 | 0 | 0 | | | | | |
| **TPJ(r)** | 0 | 0 | 0 | 0 | 0 | 0.13 | 0 | 0 | 0 | | | | |
| **mmPFC** | 0 | 0 | 0 | 0 | 0 | 0 | 0 | 0 | 0 | 0 | | | |
| **dmPFC** | 0 | 0 | 0 | 0 | 0 | 0 | 0 | 0 | 0 | 0 | 0.17 | | |
| **vmPFC** | 0 | 0 | 0 | 0 | 0 | 0 | 0 | 0 | 0 | 0 | 0 | 0 | |
